# Several cell-intrinsic effectors drive type I interferon-mediated restriction of HIV-1 in primary CD4^+^ T cells

**DOI:** 10.1101/2023.02.07.527545

**Authors:** Hannah L. Itell, Daryl Humes, Julie Overbaugh

## Abstract

Type I interferon (IFN) upregulates proteins that inhibit HIV within infected cells. Prior studies have identified IFN-stimulated genes (ISGs) that impede lab-adapted HIV in cell lines, yet the ISG(s) that mediate IFN restriction in HIV target cells, primary CD4^+^ T cells, are unknown. Here, we interrogate ISG restriction of primary HIV in CD4^+^ T cells. We performed CRISPR-knockout screens using a custom library that specifically targets ISGs expressed in CD4^+^ T cells and validated top hits. Our investigation identified new HIV-restricting ISGs (HM13, IGFBP2, LAP3) and found that two previously studied factors (IFI16, UBE2L6) are IFN effectors in T cells. Inactivation of these five ISGs in combination further diminished IFN’s protective effect against six diverse HIV strains. This work demonstrates that IFN restriction of HIV is multifaceted, resulting from several effectors functioning collectively, and establishes a primary cell ISG screening model to identify both single and combinations of HIV-restricting ISGs.

## INTRODUCTION

The type I interferon (IFN) response is one of the first defenses of the host innate immune system that viruses must overcome during transmission and acute infection. IFN signaling, triggered by viral infection, upregulates hundreds of IFN-stimulated genes (ISGs), many of which inhibit viral replication in a cell-intrinsic manner within infected cells. The ISGs that are effective at preventing viral replication vary in their magnitude of restriction as well as by virus, host, and cell type ^1^. Like other viruses, HIV-1 is sensitive to IFN restriction. Cell culture work has extensively characterized type I IFN-mediated inhibition of HIV replication ^2–6^ and studies of IFN treatment in chronically infected patients report reduced viral loads ^7–10^, demonstrating the efficacy of IFN restriction *in vivo*. Because type I IFN is secreted early after natural HIV infection ^11, 12^ and inhibits HIV replication *in vivo*, the IFN response may significantly contribute to HIV transmission barriers, transmission bottlenecks, and early viral load differences, all of which influence patient outcomes.

Due to this clinical potential, there is considerable interest in identifying the cellular ISGs that inhibit HIV replication. Numerous studies have pursued this goal and described HIV restriction mediated by ISGs such as TRIM5*α*, IFITMs, SAMHD1, SLFN11, MX2, Tetherin, APOBEC3s, and many others (reviewed in ^13–15^). This extensive body of research has informed determinants of species-specificity for HIV-1 and related retroviruses ^16–19^ and uncovered diverse mechanisms of cell-intrinsic inhibition of HIV. However, two key issues remain that limit our understanding of the proteins that are clinically relevant to IFN-mediated HIV restriction. First, the majority of this work was performed in cell lines with lab-adapted HIV strains or pseudoviruses ^20–31^, and the restrictive capacity of some ISGs has varied based on these experimental conditions ^22, 32–35^. Thus, to understand the role of these factors in HIV pathogenesis, the antiviral effects of previously described HIV-restricting ISGs need to be examined in the main cell type that supports HIV replication *in vivo*, CD4^+^ T cells, against replicating viruses that have not been lab-adapted (primary viruses) and that best represent globally circulating viruses. Second, the reported inhibition of known ISGs does not fully account for the overall magnitude of IFN restriction of HIV. This suggests that either the ISGs studied thus far exhibit more pronounced restriction in primary target cells, or additional HIV-restricting ISGs have yet to be identified.

To determine the ISGs that have relevance restricting HIV in primary target cells, we inactivated candidate ISGs via CRISPR-Cas9 editing in primary CD4^+^ T cells and infected edited cells with two diverse replication-competent HIV strains, including a primary virus and a lab-adapted strain. We first targeted eight ISGs previously studied in other infection systems but found that only one gene (IFI16) contributed to detectable IFN restriction in our model. To identify additional candidate ISGs, we performed comprehensive CRISPR-knockout (KO) screens in CD4^+^ T cells using a custom library reported here (CD4-ISG library) that targets ISGs specifically upregulated in this important cell type. Screens with the CD4-ISG library identified several candidate ISGs. Half of the eight hits we pursued further (HM13, IGFBP2, LAP3, UBE2L6) demonstrated IFN-dependent restriction of the primary HIV strain, including three factors not previously known to inhibit HIV. Because each IFN effector alone did not explain the entirety of IFN restriction, we inactivated all five ISGs in CD4^+^ T cells and found that IFN inhibition was more fully ablated as additional genes were targeted. These findings demonstrate that IFN blocks HIV by upregulating several effector proteins that work collectively. This study also describes a valuable primary CD4^+^ T cell ISG screening model for the identification of both single and combinations of antiviral ISGs, including factors that have been overlooked in culture-adapted models of HIV infection.

## RESULTS

### *Ex vivo* CD4^+^ T cell model to define ISG restriction of HIV

To define the effectors of IFN-mediated restriction of HIV in primary target cells, we interrogated ISGs of interest by inactivating them via CRISPR-Cas9 editing in human CD4^+^ T cells isolated from whole blood (**Figure 1A**). Activated CD4^+^ T cells were nucleofected with commercial triple-guide CRISPR-Cas9 ribonucleoproteins (RNPs) to achieve high levels of ISG KO. To recapitulate natural infection, we established spreading infections (MOI=0.02) with a replication-competent, CCR5-tropic primary HIV strain (Q23.BG505, clade A) ^36, 37^. To better align our results with previous studies, we also infected with a commonly used CXCR4-tropic lab-adapted HIV strain (LAI, clade B) ^38^ for comparison. IFN-*β* was chosen as the type I IFN for this model as it elicits a more robust, prolonged transcriptional response as compared to IFN-*α* ^39^. To capture the effect of ISGs on any stage of viral replication, we evaluated infection by measuring reverse transcriptase (RT) activity in the supernatant.

**Figure 1.**
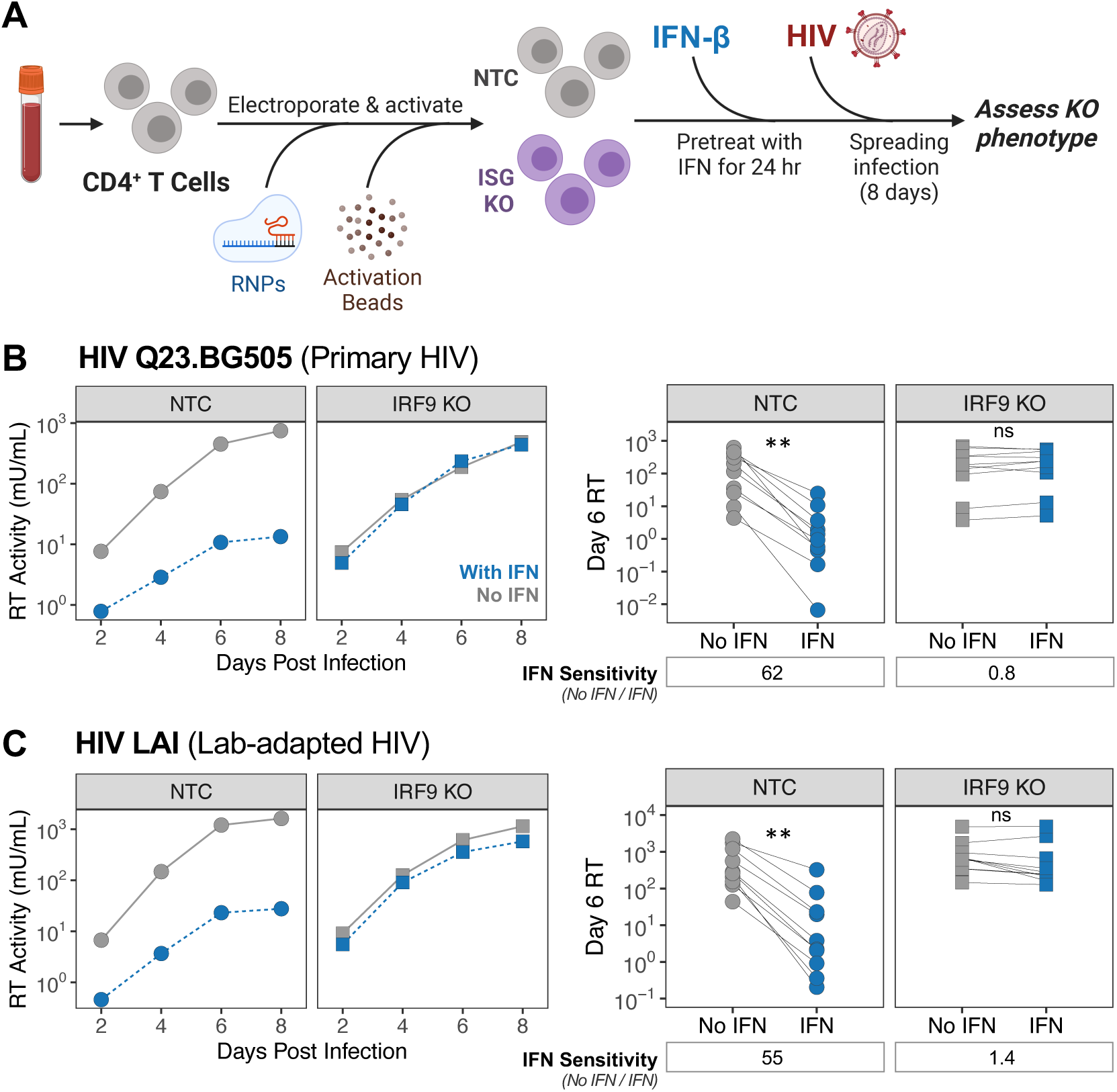
*Ex vivo* CD4^+^ T cell model to define ISG restriction of HIV. (**A**) Schematic of approach for evaluating ISG function in primary CD4^+^ T cells. (**B, C**) NTC-delivered and IRF9-edited CD4^+^ T cells were infected with full-length HIV strains Q23.BG505 and LAI, respectively, for eight days at an MOI of 0.02. Reverse transcriptase (RT) activity was measured over the course of infection. Left, representative infection curves from one donor comparing RT levels from infections with and without IFN treatment (With IFN, No IFN). Right, 6 days post infection (dpi) RT activity from infections with and without IFN from 10 independent experiments that used 8 unique donors. Dot plots are paired by experiment and the median IFN sensitivity across these experiments are depicted below (IFN sensitivity = untreated day 6 RT / with IFN day 6 RT). **p<0.002, paired Wilcoxon rank test. Non-significant (ns) indicates p>0.05. See also **Figure S1**.

In non-targeting control (NTC)-delivered CD4^+^ T cells from several independent donors, both viruses exhibited consistent sensitivity to IFN treatment (Q23.BG505 median: 62-fold; LAI median: 55-fold; **Figure 1B and 1C**). Editing of IRF9, a component of the ISGF3 transcription factor complex that enables expression of ISGs, served as a positive control and led to high KO levels for all donors (median: 94.5% KO, range: 62-100% KO; **Figure S1A**). As expected, IRF9 KO effectively blocked ISG protein upregulation as measured by Western blot analysis of the MX2 ISG (**Figure S1B**), and completely ablated the inhibitory effect of IFN for both viruses (**Figure 1B and 1C**). Of note, we did not observe a substantial increase in HIV levels with IRF9 KO in the absence of IFN treatment, which suggests that infected cells are not endogenously secreting inhibitory levels of type I IFN in this system (**Figure S1C and S1D**). This *ex vivo* model of CD4^+^ T cells infected with an IFN-sensitive primary HIV strain therefore represents a robust, biologically tractable approach for measuring the contribution of ISGs to IFN restriction of HIV.

### IFI16 contributes to IFN restriction of primary HIV in CD4^+^ T cells

We first interrogated the inhibitory function of eight proteins previously characterized as possible effectors of IFN restriction of HIV in other systems. IFI16, IFITM1, MX2, and TRIM5*α* were selected based on their ability to confer cell-intrinsic restriction of HIV in studies using lab-adapted viruses, pseudoviruses, or cell line models ^17, 20–22, 26, 30, 31, 40–42^. Of note, IFITM1 has been evaluated in primary CD4^+^ T cells in one study that found it contributed to IFN restriction of one out of two IFN-sensitive primary strains ^43^. APOBEC3A (A3A), BCL2L14/BCLG, CMPK2, and IFI27, as well as the previously selected IFITM1 and MX2, were targeted because their increased expression in CD4^+^ T cells correlated with decreased plasma viral loads in IFN-treated patients with chronic HIV infection ^9^.

Because of the inherent variability of primary cells from different donors, we performed KO experiments with 4-5 donors for each of the eight genes of interest. Nucleofection with Cas9 RNPs yielded median gene KO scores ranging from 79% (A3A) to 100% (IFI16, IFITM1, MX2), with all genes and donors exceeding 30% KO, a previously established functional cutoff for genomic editing ^44^ (**Figure 2A**). Negative control cells were IFN-treated and infected in parallel to ISG KOs to contextualize KO results. To determine whether each ISG meaningfully contributed to IFN inhibition, we applied two criteria based on IFN restriction and HIV infection phenotypes at 6 days post infection (dpi), as illustrated in **Figure S2A**. First, to assess whether ISG KO is contributing to IFN restriction, we required that ISG KO must increase HIV levels in the context of IFN treatment by at least two-fold as compared to the intra-assay negative control. We refer to this metric as “HIV Levels with IFN”. In cases where this condition was met, we applied a second criterion to eliminate genes whose inhibition is not specifically associated with IFN. For this, we calculated each KO’s relative IFN sensitivity, which is the ratio of gene KO without IFN to gene KO with IFN. We required that the IFN sensitivity of ISG KO must be at least two-fold lower than that of the negative control. We refer to this metric as “Relative IFN Sensitivity”. These criteria effectively select for genes that are IFN effector proteins as opposed to non-ISG restriction factors or genes whose KO impairs HIV infection (**Figure S2B**). For instance, KO of a non-ISG restriction factor would increase HIV Levels with IFN, but would not decrease Relative IFN Sensitivity, because infection levels would be increased by the KO independent of IFN treatment. Likewise, if gene KO hinders HIV replication, such as in cases of diminished cell growth or viability, infection levels both with and without IFN would be decreased, which may misleadingly decrease Relative IFN Sensitivity. By first requiring increased HIV Levels with IFN, this gene would not be categorized as an IFN effector. IRF9 KO appropriately meets these criteria (**Figure S1E**), causing a 160-fold increase in HIV Levels with IFN and a 78-fold decrease in Relative IFN Sensitivity.

**Figure 2.**
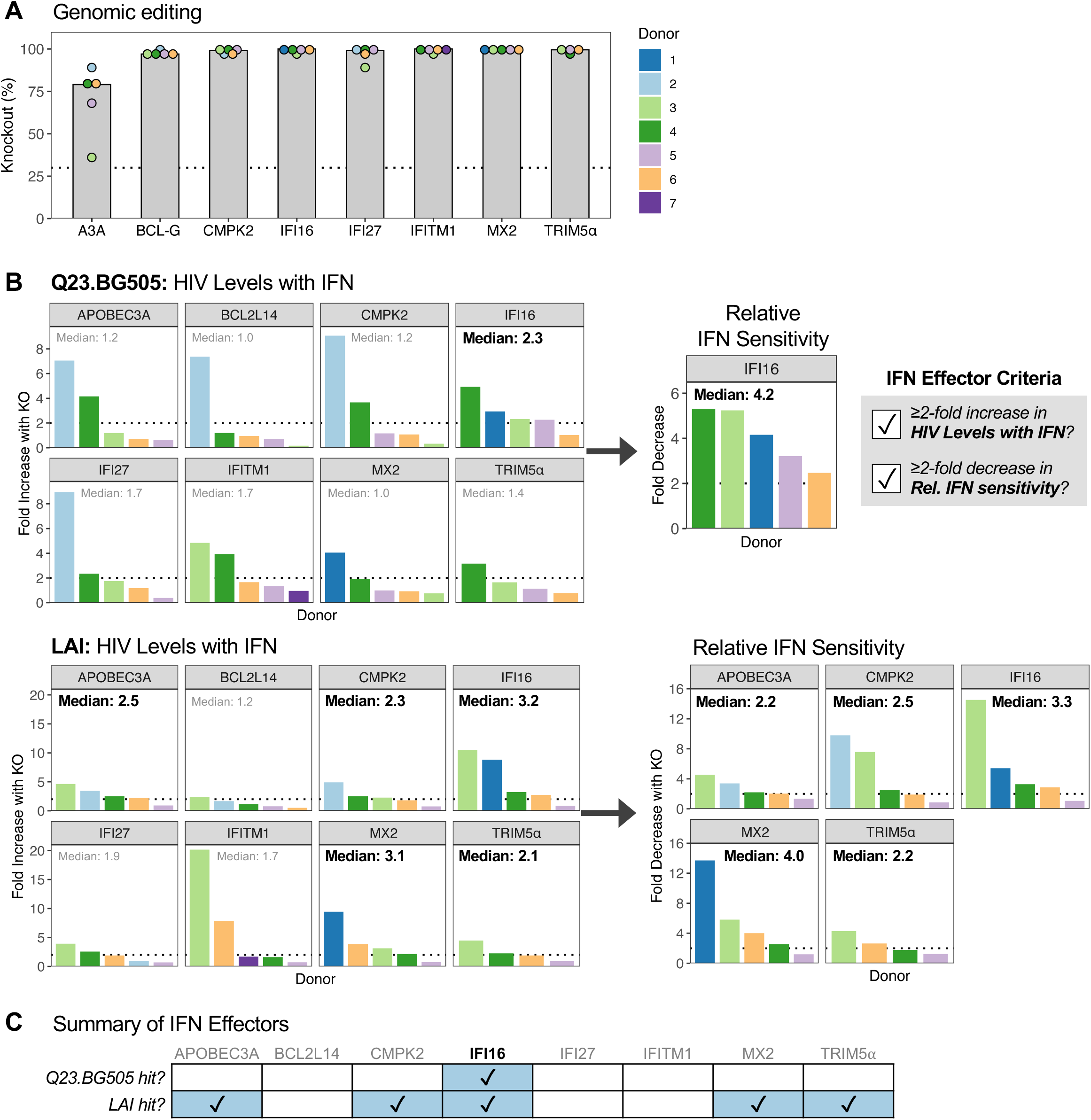
IFI16 contributes to IFN restriction of primary HIV in CD4^+^ T cells. (**A**) Nucleofected CD4^+^ T cells were collected on the day of infection and assessed for genomic editing by Sanger sequencing. Synthego ICE analysis was applied to sequencing results to determine percent knockout (KO). The median percent KO across 4-5 donors is depicted. The dotted line denotes 30% KO. Donor color is consistent within the figure but does not correspond with other figures unless otherwise noted. (**B**) ISG-edited and NTC- delivered cells were treated with or without IFN and infected with Q23.BG505 (top) and LAI (bottom). Left, increase in HIV Levels with IFN (6 dpi RT levels) with ISG KO as compared to NTC cells (ISG KO/NTC). The median increase across donors is depicted at the top of each graph. The dotted line indicates a fold difference of two. Right, for genes on the left with median increases in HIV Levels with IFN ≥2-fold, the decrease in Relative IFN Sensitivity with ISG KO compared to NTC is shown (NTC/ISG KO). (**C**) Summary of ISG KO infection phenotypes. See also Figures S2 and S3.

As expected for primary CD4^+^ T cells, we observed some variability in infection phenotypes between donors, such as A3A, BCL2L14, CMPK2, and IFI27 KOs demonstrating much stronger impacts on Q23.BG505 HIV Levels with IFN in Donor 2 as compared to the four other donors (**Figure 2B**). We therefore evaluated median fold effects of the 4-5 donors tested by our two analysis criteria to select ISGs with restrictive activity across donors. Surprisingly, only IFI16 met both criteria for the primary HIV strain Q23.BG505, with knockout causing a median 2.3-fold increase in HIV Levels with IFN, which corresponded to a median 4.2-fold decrease in Relative IFN Sensitivity (**Figure 2B and 2C**). IFI16, as well as A3A, CMPK2, MX2, and TRIM5*α*, met both criteria for LAI. MX2 KO had the largest impact on the Relative IFN Sensitivity for this virus (median 4-fold decrease). These data define IFI16 as a broad effector of IFN restriction of HIV and suggest that other tested ISGs, most notably MX2, IFITM1, and TRIM5*α*, have less relevance in primary target cells against a primary virus than in the models in which they were previously described.

Because MX2 has been shown to restrict several HIV strains in various cell lines ^21, 26, 30, 31, 35, 45–48^, we were surprised that editing of MX2 did not meaningfully increase Q23.BG505 HIV Levels with IFN, like it did for LAI (**Figure S3A**). The HIV Capsid (CA) is the main viral determinant of MX2 restriction. We therefore aligned the CA sequence of Q23-17 ^36^, which encodes the *gag* gene in Q23.BG505, to those of other MX2-resistant strains (RHPA.c and CH040.c) and MX2-sensitive strains (LAI/IIIB, NL4-3, CH058.c, CH077.t, CH106.c, YU-2 (CA identical to REJO.c), and WITO.c) (**Figure S3B and S3C**). This identified 18 residues in which Q23-17 differed from all MX2-sensitive CAs, including four residues located in regions where mutations are known to confer MX2 escape: V11I, E71D, A92P, E187D. Capsid differences between LAI and Q23.BG505 may therefore contribute to their differential sensitivity to MX2. Interestingly, these positions are also variable across HIV clades, with the D71 and P92 residues found in Q23-17 representing the Group M consensus sequence (**Figure S3D**).

### Custom CRISPR sgRNA library to target ISGs in CD4^+^ T cells

Though IFI16 KO reduced the Relative IFN Sensitivity of Q23.BG505 (median 4.2-fold, **Figure 2B**), it only captured a fraction of the effect seen with IRF9 KO (median 78-fold, **Figure S1E**). This discrepancy indicates that other ISGs must contribute to IFN inhibition. To evaluate the restrictive capacity of all possible ISGs expressed in primary CD4^+^ T cells, we sought to carry out comprehensive, high-throughput CRISPR-KO screens with a customized CRISPR sgRNA library relevant to these cells. We therefore characterized ISG expression in our *ex vivo* model by performing bulk RNA-seq on primary CD4^+^ T cells collected 0, 3, 6, 12, and 24 hours post IFN treatment (**Figure 3A, Table S1**). Principle component analysis revealed clear separation between IFN-treated and untreated cells and appreciable differences between collection time points (**Figure 3B**). Due to the differences between time points, we assessed sampling times individually. For results from 6, 12, and 24 hours post-IFN, we selected genes that were IFN-stimulated by at least two-fold in the majority of donors for a given time point (**Figure 3C**). Because we only had one donor with cells collected at 3 hours post-IFN, we applied a more stringent cutoff of four-fold induction for this sample. This approach yielded 261, 449, 261, and 167 genes from the 3-, 6-, 12-, and 24-hour time points, respectively, with many ISGs overlapping across sampling times (**Figure 3D**). To create the sgRNA library, we designed eight guides per gene and included 200 NTC guides as controls. The final “CD4-ISG” library targets 555 ISGs represented across 4,579 guides (**Table S2**). We compared this library to a larger, less focused library used to identify HIV antiviral ISGs in other studies (PIKA^22^) and found that the majority of genes (57%) were unique to the CD4-ISG library (**Figure 3E**). Likewise, only 12.5% of the PIKA library represents genes that are IFN-stimulated in primary T cells. The CD4-ISG library therefore encompasses a unique set of ISGs tailored specifically to primary CD4^+^ T cells.

**Figure 3.**
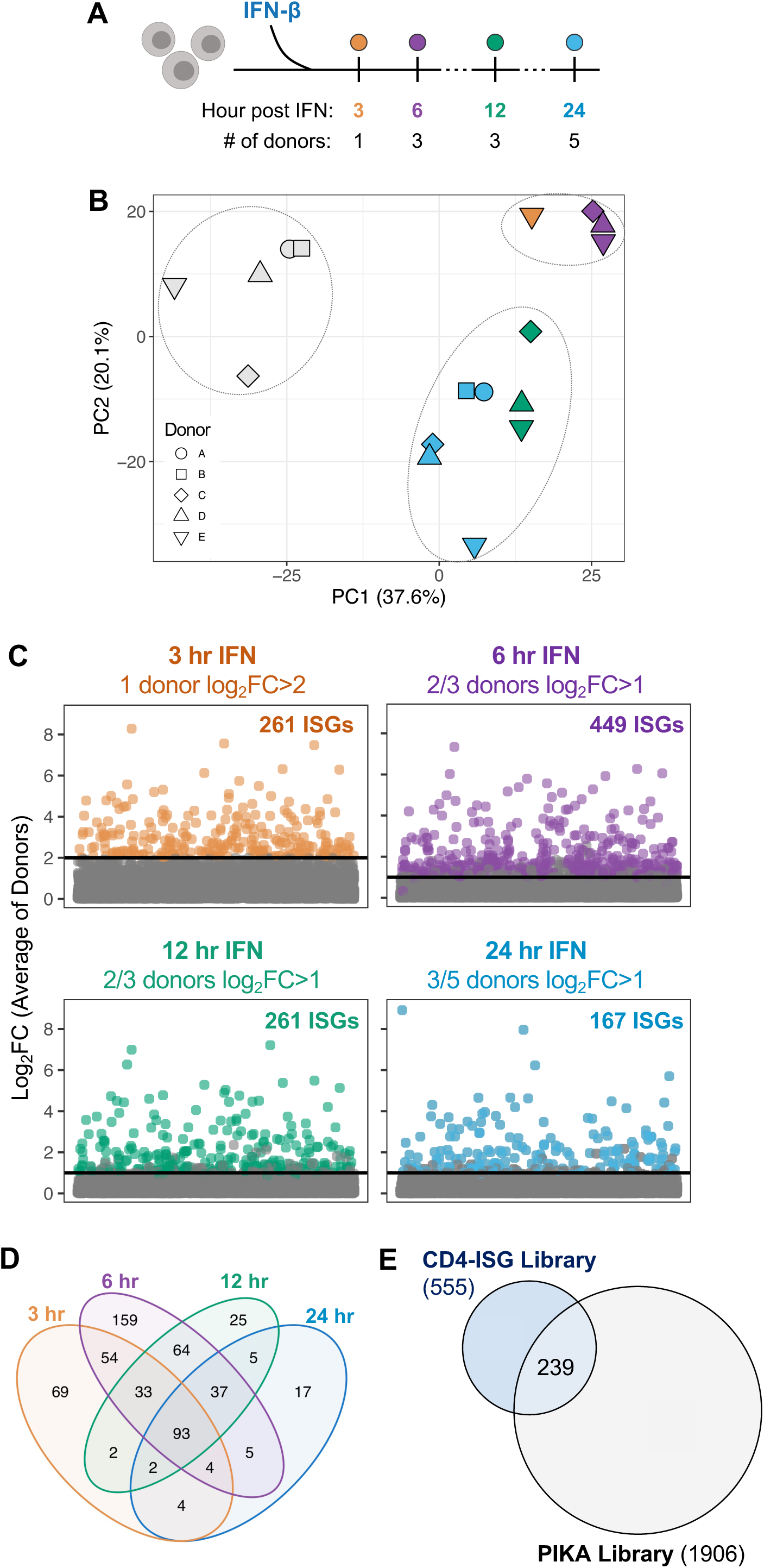
Custom CRISPR sgRNA library to target ISGs in CD4^+^ T cells. (A) Schematic describing CD4^+^ T cells evaluated by bulk RNA-seq. (**B**) Raw RNA-seq counts were corrected for donor differences using ComBat-Seq and data were visualized by principal component analysis. (**C**) To select ISGs to include in a new CRISPR library, thresholds were applied to each RNAseq time point, as indicated by the solid line. Genes above this threshold are shown in color and total ISG counts are included in each plot. (**D**) Venn diagram comparing ISGs selected in (C) by collection time. (**E**) Euler diagram comparing genes represented in the CD4-ISG libraries to those in the broad PIKA ISG library (OhAinle *et al*., 2018).

### Strategy for HIV-CRISPR screens in primary CD4^+^ T cells

To perform CRISPR-KO screens with the CD4-ISG library in primary CD4^+^ T cells (**Figure 4A**), we leveraged the HIV-CRISPR approach, a robust strategy developed in the THP-1 monocytic cell line and reported in OhAinle *et al* ^22^. HIV-CRISPR screens deliver sgRNA libraries to cells using a version of a lentiviral vector that has a repaired, complete HIV Long-Terminal Repeat (LTR). This repair allows the integrated vector, including the sgRNA sequence, to be transcribed and packaged into budding virions during replication-competent HIV infection. Therefore, guide sequences enriched in the supernatant viral RNA as compared to the cellular genomic DNA would be expected to target and inactivate genes that restrict HIV.

**Figure 4.**
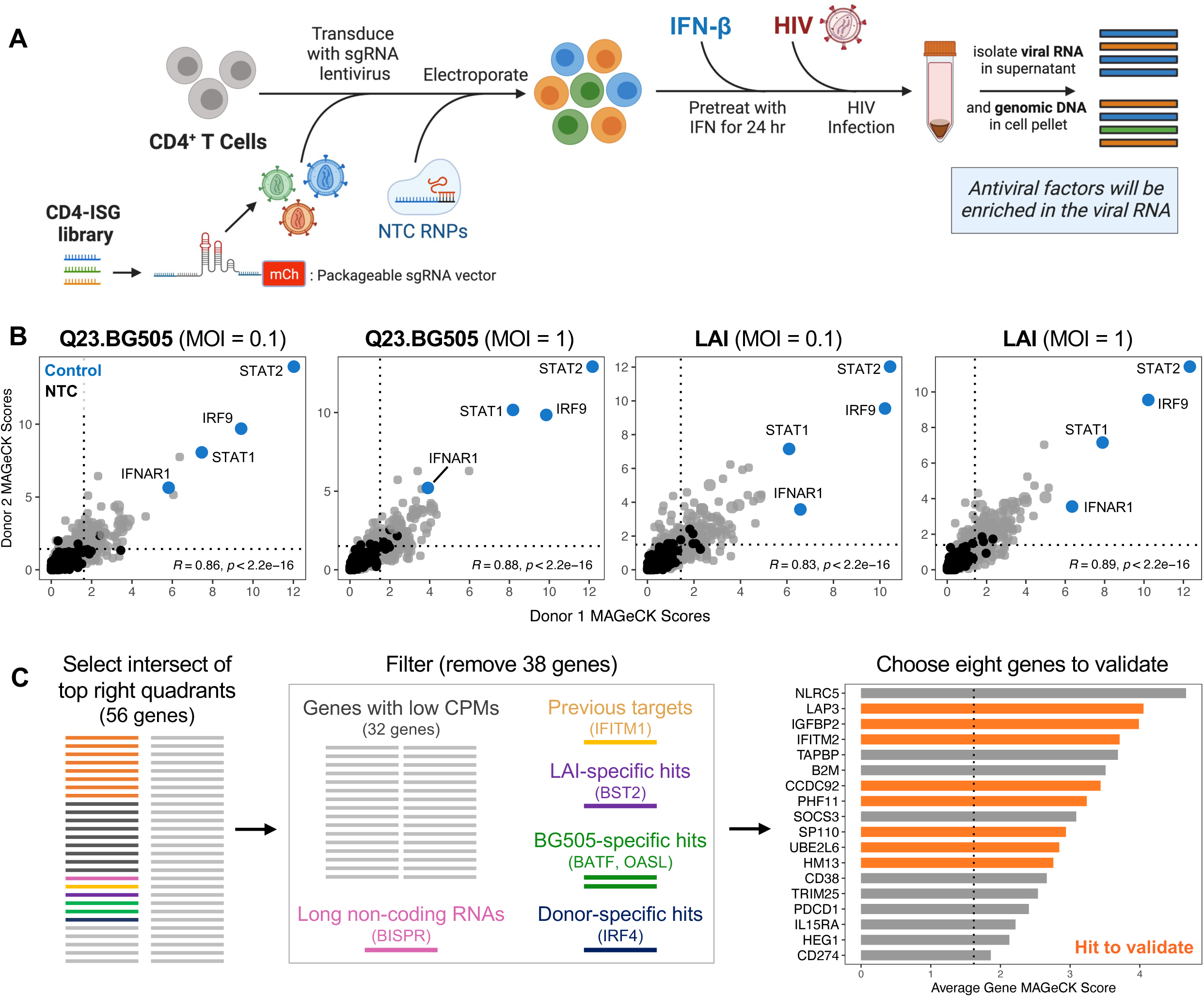
HIV-CRISPR screens with the CD4-ISG library identify HIV-restricting ISGs. (**A**) Schematic of HIV-CRISPR screens to identify HIV-restricting ISGs in primary CD4^+^ T cells. (**B**) HIV-CRISPR screens were conducted in CD4^+^ T cells from two donors. Gene-level MAGeCK enrichment scores were correlated between donors for each virus and MOI. Dotted lines indicate the NTC average + 3 standard deviations for individual screens. Pearson correlation coefficients are included within plots. (**C**) Strategy for selecting eight CD4-ISG screen hits to validate. Right, the average MAGeCK score across all eight screens for the top 18 screen hits. Dotted line denotes the highest background threshold from panel B plots. See also **Figure S4**.

CRISPR screening in primary CD4^+^ T cells presents three challenges that we had to overcome. First, lentiviral delivery of sgRNAs to CD4^+^ T cells has been prohibitively inefficient in the past. To circumvent this issue, we utilized the TOP vector, which yields high-titer lentivirus with improved transduction of primary T cells and increased packageability as compared to the widely used lentiGuide construct ^49^. To amend the TOP vector for HIV-CRISPR screening in primary T cells, the HIV LTR was repaired to restore packaging. The tracrRNA sequence was also optimized for more efficient sgRNA expression ^50^, an adjustment that was essential to achieve sufficient editing (data not shown). Second, lentiviral delivery of Cas9 is insufficient for high levels of editing in primary T cells ^51^. We adopted a previously utilized strategy that delivers Cas9 to primary T cells by nucleofection of NTC Cas9 RNPs after lentiviral transduction with the guide library ^52^. Lastly, most CRISPR screens transduce cells at low MOIs followed by antibiotic selection over several days to generate a pool of cells containing one sgRNA. Because primary T cells have short *ex vivo* culture windows, antibiotic selection is not practical. We therefore transduced cells at a high MOI, aiming to deliver roughly 3 vector copies per cell to maximize the fraction of cells expressing sgRNAs. These modifications enabled us to perform HIV-CRISPR screens in primary CD4^+^ T cells.

### HIV-CRISPR screens with the CD4-ISG library identify HIV-restricting ISGs

We conducted HIV-CRISPR screens with the custom CD4-ISG library in CD4^+^ T cells from two independent donors (**Figure 4A**). Lentiviral transductions delivered 3.0 and 5.2 vector copies per cell for each donor, respectively, and library coverage was maintained (>1000-fold) at every stage. We infected transduced cells with HIV at higher MOIs (0.1, 1) in screens than in validation experiments (MOI=0.02) to ensure sufficient read out. Genomic DNA and viral RNA deep sequencing was analyzed by MAGeCK-Flute ^53^ (**Table S3**) to quantify gene-level results and identify guides enriched in the supernatant, which would reflect the inactivation of restriction factors. Positive control IFN signaling genes (IRF9, STAT1, STAT2, IFNAR1) were highly enriched in all screens and we observed strong agreement between donors (Pearson R: 0.83-0.89; **Figure 4B**). A high proportion of genes (20.5-30%) scored above background for both donors, with LAI screens yielding slightly more hits (Q23.BG505 MOI 0.1: 114/555; Q23.BG505 MOI 1: 134/555; LAI MOI 0.1: 140/555; LAI MOI 1: 166/555). Overall, 56 ISGs scored above background in all eight screens.

We next aimed to validate genes with the highest likelihood of having broad restrictive activity in primary CD4^+^ T cells (**Figure 4C**). To select this subset, we rationalized that genes with consistent robust expression across donors would be less likely to arise as donor-specific artifacts. Therefore, from the list of 56 top-scoring genes (**Table S4**), we filtered out those with average counts per million (CPMs) across donors less than 20. We also removed one long non-coding RNA, as they are often not amenable with CRISPR approaches, and one ISG that we had previously interrogated (IFITM1, **Figure 2**). From the remaining candidates, we identified and removed strain- or donor-specific hits: LAI-specific BST2 (Tetherin), Q23.BG505-specific BATF and OASL, and donor-specific IRF4 (**Figure S4**). We selected eight of the remaining genes to validate.

ISG editing exceeded 30% KO for all genes (**Figure 5A**) and did not impact cell growth (**Figure 5B**), indicating that these ISGs did not artificially arise as screen hits due to their inactivation causing cell death. Compared to the negative control, inactivation of HM13, IGFBP2, LAP3, and UBE2L6 increased Q23.BG505 HIV Levels with IFN ≥2-fold, which corresponded with ≥2-fold decreased Relative IFN sensitivity, indicating that they are IFN effectors (**Figure 5C**). Editing of HM13 and UBE2L6 also met both IFN effector criteria for LAI, suggesting that these genes may broadly restrict diverse HIV strains. Though IFITM2 KO did not increase Q23.BG505 HIV Levels with IFN, it did demonstrate IFN effector activity against LAI. Ultimately, half of the CD4- ISG screen hits we interrogated (4/8) validated as mediators of IFN restriction of a primary virus (**Figure 5D**).

**Figure 5.**
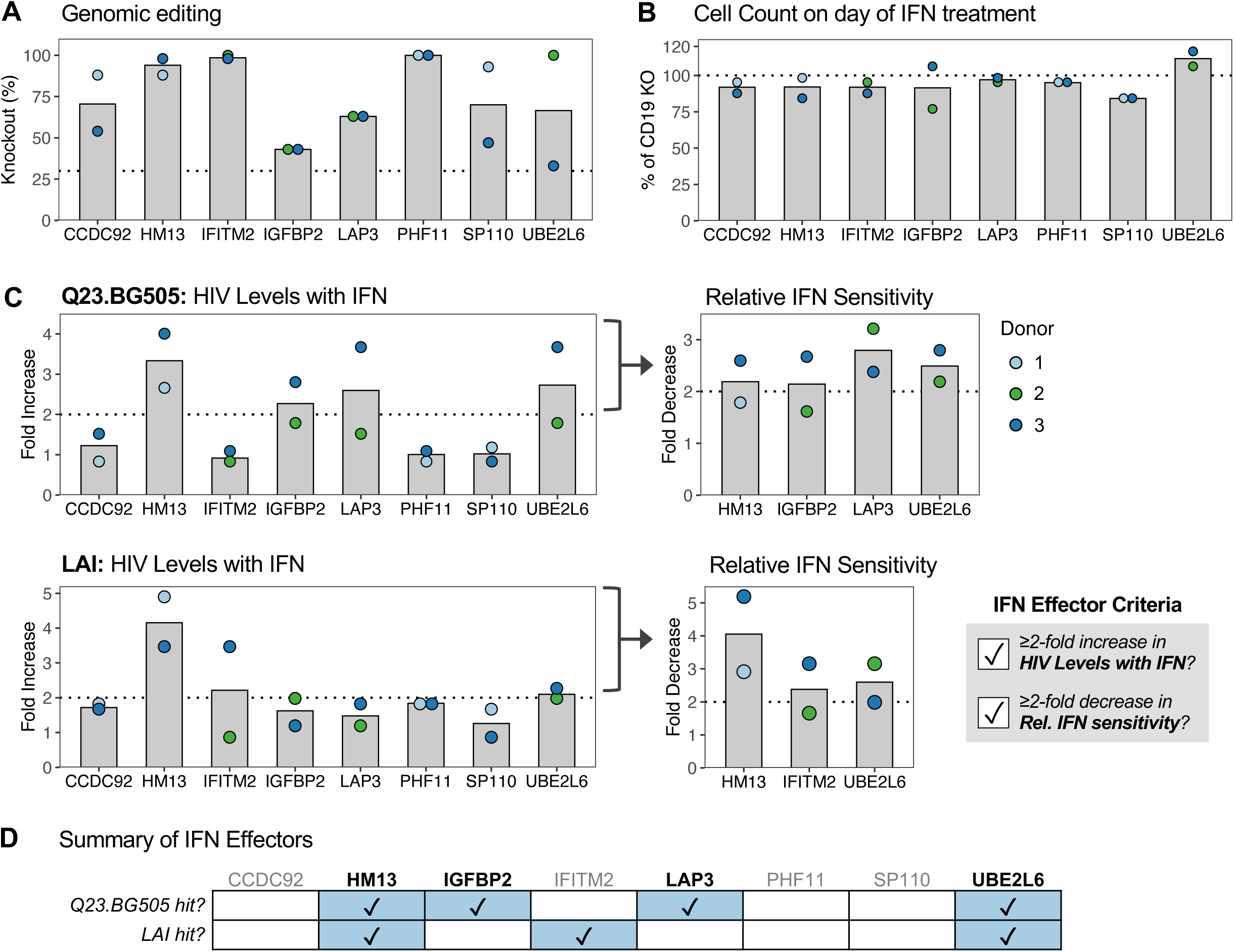
CD4-ISG HIV-CRISPR screen hits validate as IFN effectors. ISGs from Figure 4C (in orange) were validated by individually inactivating each gene in CD4^+^ T cells from two independent donors. (**A**) ICE KO percentages. Dotted line indicates 30% KO. (**B**) The number of cells for each ISG KO on the day of IFN treatment are shown relative to CD19-edited, negative control cells. Dotted line at 100%. (**C**) Left, the increase in HIV Levels with IFN with ISG KO compared to CD19 KO cells (ISG KO/CD19 KO). Right, for ISG KOs that exceeded a two-fold increase in HIV Levels with IFN, the decrease in Relative IFN Sensitivity is depicted (CD19 KO/ISG KO). (**D**) Summary of ISG KO infection phenotypes.

### IFN restriction of HIV is driven by a combination of IFN effectors

We have thus far identified HM13, IFI16, and UBE2L6 as mediators of IFN restriction of both tested HIV viruses. Additionally, IGFBP2 and LAP3 demonstrate IFN-dependent inhibition of the primary HIV strain Q23.BG505. However, the effect of each single knockout for both viruses (**Figures 2B and 5C**) is less substantial that the entire restrictive effect of IFN (**Figure 1**). We therefore hypothesized that IFN inhibition of HIV results from several IFN effectors functioning collectively. To test this hypothesis, we performed multi-gene knockouts, targeting two (HM13, IFI16), three (HM13, IFI16, UBE2L6), or all five ISGs (HM13, IFI16, UBE2L6, IGFPB2, LAP3) within the same cell pool. To extend this study and address whether these factors inhibit diverse HIV strains, we infected cells with a broader panel of six IFN-sensitive viruses, including both primary-isolated and lab-adapted viruses and viruses from different HIV clades (**Figure 6A**).

**Figure 6.**
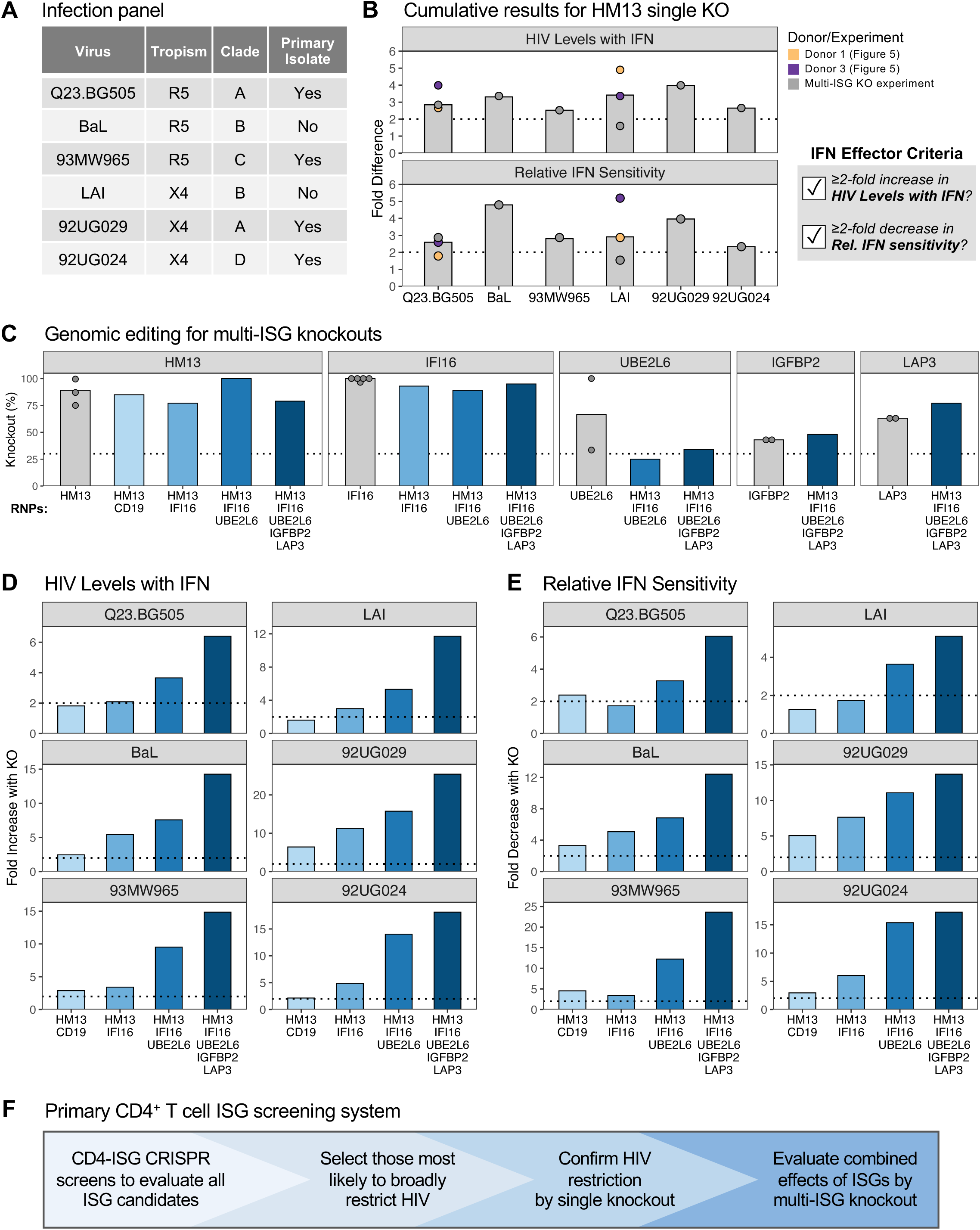
IFN restriction of HIV is driven by a combination of IFN effectors. CD4^+^ T cells from one donor were nucleofected with one to five RNPs to assess the effect of inactivating several ISGs in combination. (**A**) Characteristics of the HIV viruses used for infections. (**B**) The fold difference in HIV Levels with IFN (HM13 KO/CD19 KO) and Relative IFN Sensitivities (CD19 KO/HM13 KO) between HM13 single KO and CD19 single KO cells for each virus tested. Dotted lines indicate a fold difference of two. (**C**) Percent KO in cells nucleofected with single (gray) or multiple (blue) RNPs, as denoted along the x-axis. Results are separated by editing locus, as indicated above each panel. Single KO bars include results from previous experiments with donors represented as points (n=2-5). The dotted line denotes 30% KO. (**D**) The increase in HIV Levels with IFN (ISG KO/CD19 KO) and (**E**) the decrease in Relative IFN Sensitivities (CD19 KO/ISG KO) for each multi-ISG KO compared to CD19 KO, for all viruses tested. Dotted lines indicate a fold difference of two. (**F**) Schematic of the screening model established in this report. See also **Figure S5**.

We selected HM13 as the intra-experiment single ISG KO control, as HM13 editing led to the greatest increase in HIV Levels with IFN for both viruses (Q23.BG505: 3.3-fold; LAI: 4.2-fold). Like we previously observed with Q23.BG505 and LAI, HM13 met both IFN effector criteria for all additional HIV strains we tested (BaL, 93MW965, 92UG029, 92UG024), with its KO leading to an increase in HIV Levels with IFN and a decrease in Relative IFN Sensitivity that was generally similar across viruses (HIV Levels with IFN: 2.5- to 4-fold, Relative IFN Sensitivity: 2.3- to 4.8-fold; **Figure 6B**). These data confirm that HM13 broadly contributes to IFN restriction of a diverse set of HIV strains and is therefore a stringent single ISG KO control.

Multi-gene knockouts were achieved by nucleofecting primary CD4^+^ T cells with RNPs targeting multiple ISGs. This strategy was successful, yielding editing efficiencies similar to prior respective single-gene KOs (**Figure 6C**). Though UBE2L6 editing in the triple KO fell below the 30% threshold (25% KO), this level of inactivation is similar to one of the previous UBE2L6 single-gene KOs (33%, Donor 3 in **Figure 5**), which impacted HIV Levels with IFN and Relative IFN Sensitivity to similar magnitudes as Donor 2 despite having lower levels of editing. Notably, editing of up to three ISGs in one cell pool did not impact cell viability (**Figure S5A**). We did however observe a 34% reduction in the number of cells in the five-ISG KO condition, similar to a prior report that tested a six-gene KO in CD4^+^ T cells ^54^. To assess whether multi-gene editing influences HIV infection or IFN restriction non-specifically, we compared the increase in HIV Levels with IFN for HM13 single KO cells to that of cells edited for both HM13 and the negative control B cell marker CD19 (**Figure S5B**). Double KO cells closely recapitulated the infection phenotype observed in HM13 single KO cells, and there was no trend in terms of which KO caused greater effects on HIV Levels with IFN. Therefore, inactivation of two or three ISGs in the same cell pool does not impact cell viability, HIV infection, nor IFN restriction non-specifically. Moreover, it is unlikely that the lower cell counts with five-ISG KO would influence whether this condition is classified as an IFN effector or not, as our analysis criteria account for such situations (**Figure S2B**).

We next evaluated whether inactivation of several ISGs impacts IFN restriction of HIV more potently than HM13 KO alone. We observed that with each additional ISG inactivated, HIV Levels with IFN increased (**Figure 6D**) and Relative IFN Sensitivities decreased (**Figure 6E**). This trend was consistent for all six viruses, indicating that the ISGs identified in this report represent broad IFN effectors. Though the multi-ISG KOs do not fully account for IFN’s entire restrictive effect, these data provide evidence that IFN restriction of HIV is mediated by several individual antiviral ISGs functioning collectively.

## DISCUSSION

In this study, we shed light on the multifaceted nature of cell-intrinsic IFN restriction of HIV and identify new effector ISGs with likely relevance *in vivo* by establishing and employing a primary CD4^+^ T cell ISG screening system. This strategy involves performing comprehensive CD4-ISG CRISPR screens to assess all ISG candidates, selecting and validating the factors most likely to have broad antiviral activity, and evaluating ISG synergy by inactivating several genes in combination (**Figure 6F**). We found that HM13, IFI16, IGFBP2, LAP3, and UBE2L6 each partially mediate IFN-dependent restriction of HIV in primary CD4^+^ T cells. Most of these factors (HM13, IGFBP2, LAP3) were not previously known to inhibit HIV, underscoring the advantage of leveraging unbiased CRISPR screens in primary target cells for ISG candidate selection. Inactivation of the five ISGs we identified in combination demonstrated that these factors collectively contribute to the antiviral effect of IFN for a diverse panel of HIV strains, including several primary isolates. This report, to our knowledge, is the first to demonstrate that the inhibitory effect of IFN is the result of several antiviral ISGs working in concert and the first to comprehensively examine ISGs that inhibit primary HIV in primary target cells.

Of the five ISGs we found to mediate IFN restriction of HIV, most (HM13, IGFBP2, and LAP3) have not been previously recognized as having anti-HIV activity and thus represent newly identified antiviral effectors. HM13 is of particular interest as it is known to directly interact with HIV gp160 ^55, 56^ and, in this study, demonstrated broad antiviral activity against all tested HIV strains both individually and in combination with other ISGs. HM13, also referred to as signal peptide peptidase (SPP), cleaves signal peptides in the endoplasmic reticulum after they have been released from polyproteins ^57^, preproteins, and misfolded membrane proteins ^58^. It has been implicated as a host dependency factor for HCV ^59^ and HSV-1 ^60, 61^ by mediating HCV core protein processing and binding to HSV glycoprotein K, respectively. Moreover, it may have dual roles both promoting and inhibiting DENV replication at different stages ^62, 63^. HM13 therefore interacts with several other viruses, and, in the case of HCV, this interaction relies on HM13’s peptidase activity. Collectively, these findings suggest that future studies should investigate whether HM13 cleaves HIV gp160 to inhibit viral replication.

Unlike HM13, the potential function of LAP3 and IGFBP2 during viral replication is less clear. LAP3 is a aminopeptidase primarily located in the cytosol that catalyzes the removal of N-terminal hydrophobic amino acids from various peptides and proteins ^64^. IGFBP2 is known for binding insulin-like growth factors when secreted but it has many additional intracellular ligands and intrinsic roles, typically associated with cell survival ^65^. Interestingly, IGFBP2 was the second highest-scoring hit in a study examining HIV Vpu-mediated suppression of cytokines, suggesting that its expression may impede viral replication ^66^. Because LAP3 and IGFBP2 have not been studied in the context of infection previously, additional work is required to elucidate their mechanism as HIV restriction factors.

In addition to identifying new antiviral effector proteins, our findings indicate that IFI16 and UBE2L6, which have been previously studied in other HIV infection models, mediate IFN restriction of several primary HIV isolates in primary CD4^+^ T cells. IFI16 has two known mechanisms of HIV restriction – it can function as a cytosolic immune sensor of HIV DNA intermediates, which results in IFN secretion and pyroptosis ^67–69^ to indirectly block HIV replication, and it sequesters the transcription factor Sp1 to broadly inhibit retroviral transcription ^20^. If IFI16 was impacting HIV infection in our model via sensing and IFN production, we would expect to see increased levels of HIV with IFI16 KO in the absence of exogenous IFN treatment. Instead, we observed minimal effects, with IFI16 KO causing a median 1.4- and 1.0-fold decrease in HIV without IFN for Q23.BG505 and LAI, respectively. Therefore, our findings are likely the result of IFI16 directly restricting HIV by reducing the availability of Sp1. This mechanism was originally characterized in macrophages, which also support HIV replication both *in vitro* and *in vivo*; thus, our findings support that IFI16 also directly inhibits HIV infection in primary CD4^+^ T cells. Like IFI16, the mechanism of UBE2L6 restriction has been previous described. UBE2L6 conjugates ubiquitin (ubiquitination) or the highly IFN-inducible ISG15 (ISGylation) to other proteins, leading to broad biological impacts ^70^ including preventing HIV budding and egress ^71^. HIV Vpu targets UBE2L6 for degradation to impair ISGylation, but this antagonism is incomplete as evidenced by residual UBE2L6 protein expression and restrictive activity in THP-1 cells ^22, 72^. Our data support that Vpu-mediated antagonism of UBE2L6 is incomplete and UBE2L6 contributes to IFN restriction of replication-competent virus in primary target cells.

While HM13, IFI16, IGFBP2, LAP3, and UBE2L6 have consistent antiviral activity across several HIV strains, our results also highlight cell type- and virus-specific differences for other ISGs. For instance, IFITM1 KO did not meet our criteria as an antiviral ISG for either virus in primary CD4^+^ T cells, despite its restrictive effect in SupT1 ^40^ and U87 ^43^ cell line models. In the case of HM13, IGFBP2, and LAP3, which have not been described as HIV inhibitors in cell lines, it remains to be seen whether they are functional in other cell types. The comprehensive nature of this screen, including the use of the tailored CD4-ISG library, may well reveal antiviral factors that are active in different cell types.

Our findings also shed light on differences in the antiviral effects of ISGs between HIV strains, which we were able to capture by infecting with two viruses for all experiments: Q23.BG505 (clade A, CCR5-tropic, primary) and LAI (clade B, CXCR4-tropic, lab-adapted). Most notably, we observed LAI-specific restriction for IFITM2, MX2, and TRIM5*α*. This result for IFITM2 is consistent with the existing literature, as IFITM2 has been shown to preferentially restrict CXCR4-tropic viruses ^43, 73^. We were particularly surprised that MX2 KO did not impact Q23.BG505 replication, as MX2 has been shown to mediate IFN restriction of several HIV strains in cell lines ^21, 26, 30, 31, 35, 45–48^ and was functional against LAI in our study. This led us to identify 18 amino acids in HIV CA, which is the main viral determinant of MX2 sensitivity, that differ between Q23-17 and eight MX2-sensitive viruses. Four of the mutations reside in CA sites or regions known to influence MX2 restriction and thus are most likely to be responsible for Q23.BG505’s resistance to MX2: V11I, E71D, A92P, E187D. Interestingly, these Q23- 17 CA residues are present in many circulating viruses. D71 and P92 are utilized in the group M consensus sequence, as is I11 for clade A and D187 for clade C ^74^. Until the use of clade A Q23.BG505 in this report, all MX2 studies have solely tested clade B viruses, suggesting the possibility that MX2 may primarily target clade B viruses *in vivo*. Our findings thus indicate that to best understand the impact of MX2, future studies should include additional strains from the globally dominant clades to investigate whether MX2 restriction is clade-specific in primary CD4^+^ T cells. Given that TRIM5*α* also interacts with the HIV CA, it is possible that CA differences between Q23-17 and LAI may also contribute to Q23.BG505’s resistance to TRIM5*α* restriction.

While our approach identified five IFN effectors with broad antiviral activity, we may have only scratched the surface on the ISGs that contribute to IFN-mediated inhibition of HIV due to the stringency of our analysis and the scope of our investigation. For instance, each CD4-ISG HIV-CRISPR screen enriched for over 100 ISGs (range: 114-166 hits), which we narrowed down to 17 candidates based on their consistency of enrichment in screens and RNA expression across donors. We then selected 8 of the highest-scoring hits to validate. In fact, IFI16 did not meet our filtering criteria despite it scoring above background in half of the screens and being a validated IFN effector. Therefore, this screen identified many more ISGs that have yet to be tested, including virus-specific hits that were not pursued in this study. These data also suggest that our criteria for defining an ISG as an IFN effector was highly stringent because two genes (IGFBP2, LAP3) that met our criteria as Q23.BG505 IFN effectors did not do so for LAI (average increase in HIV Levels with IFN: 1.6-fold for IGFBP2 KO and 1.5-fold for LAP3 KO), yet multi-gene KO experiments demonstrated that editing of these factors in addition to HM13, IFI16, and UBE2L6 further decreased IFN restriction of LAI. The stringency of our IFN effector classification criteria may have therefore excluded “real” HIV-restricting ISGs. Moving forward, inactivating factors alone as well as in combination may provide the most robust data to determine whether ISGs meaningfully contribute to IFN restriction.

Though we aimed to maximize biological relevance and experimental robustness, our approach has important caveats to consider. First, primary cell models do not have unlimited cell availability. This made it impossible to use the same blood donors for all ISG KO experiments, which could contribute to experimental variation. However, we did not observe notable patterns in editing or KO phenotypes for specific donors and we controlled for differences in HIV infection levels across donors by determining the within-assay fold effect of ISG KO relative to negative controls. This allowed for direct, quantitative comparisons across independent experiments and ISG targets. Second, our single KO approach with triple-guide RNPs was relatively low throughput compared to nucleofections with arrayed single-guide RNPs ^44, 75, 76^. Though our approach limited the number of donors and viruses we could test as well as the number of screen hits we could validate, it also had valuable tradeoffs. Smaller scale nucleofections allowed us to closely monitor cell counts to ensure all KOs were accurately infected at a low MOI of 0.02. Moreover, we achieved reliable sequencing and high KO scores for all sixteen single ISG targets, whereas a recent arrayed CRISPR screen was unable to assess 18% (77/436) of targets due to incomplete sequencing information or insufficient editing ^44^.

Type I IFN upregulates hundreds of ISGs; thus, it is not surprising that we found IFN-mediated inhibition of HIV to result from many ISG effectors as opposed to one extremely potent ISG. While nearly all prior reports have inferred this conclusion due to their ISG of interest not accounting for full IFN restriction, particularly a study based on overexpressed pairs of ISGs prior to HIV pseudovirus challenge in a cell line ^77^, none to our knowledge have inactivated multiple ISGs within the same cell pool to directly support this hypothesis. Our report therefore establishes that many cell-intrinsic ISGs contribute to IFN restriction of replication-competent HIV in primary target cells, CD4^+^ T cells. Some of these effectors are genes targeted by HIV accessory proteins (UBE2L6, IGFBP2), suggesting that their antagonism is incomplete. Moreover, viral sensitivity to ISGs can vary by clade and tropism/co-receptor use (MX2, IFITM2, TRIM5*α*), which is emphasized by the findings presented here. This underscores the need to test diverse HIV strains when assessing ISG inhibitory potencies and to consider the viruses tested when extrapolating *in vitro* studies to *in vivo* mechanisms.

There is now considerable interest in identifying the minimal network of ISGs needed to ablate the majority of IFN restriction and it remains to be determined whether a potent pan-HIV ISG network exists or if ISG sets would need to be clade-specific. Leveraging the primary CD4^+^ T cell ISG screening system reported here provides the opportunity to identify additional HIV restriction factors with relevance in the cells that are responsible for the majority of HIV produced *in vivo*. In addition to the hits from our screens that have yet to be validated, future iterations of this system could perform HIV-CRISPR screens with the CD4-ISG library with different HIV strains or in the background of an ISG KO to select for ISGs with synergistic effects. Resolving the group of ISGs that account for IFN-mediated protection against HIV in primary CD4^+^ T cells would directly inform the cell-intrinsic barriers that HIV must overcome during transmission and acute infection.

## ACKNOWLEDGMENTS

We thank the Fred Hutch Shared Resources Genomics and Bioinformatics Cores, particularly Alyssa Dawson, Elizabeth Jensen, Matthew Fitzgibbon, and Pritha Chanana, for their contributions performing Sanger sequencing, Tapestation, and Illumina sequencing and assisting with bulk RNA-seq analysis and the CD4-ISG library guide design; the Fred Hutch Center for Data Visualization, especially Michael Zager, Nathan Thorpe, and Sam Minot, for their support in performing MAGeCK analysis for CD4-ISG screens; Michael Emerman, Molly OhAinle, Emily Hsieh, Vanessa Montoya, Caitlin Stoddard, Joshua Marceau, Nell Baumgarten, and Alex Willcox for helpful discussions and technical assistance. This work was supported by the following NIH grants: NICHD R01 HD103571 and NIDA DP1 DA039543 to J.O and NIGMS T32 GM007270 and NIAID F31 AI165168 to H.L.I.

## AUTHOR CONTRIBUTIONS

J.O. conceived the project and H.L.I., D.H., and J.O. designed the methodology of the study. H.L.I. and D.H. performed experiments. H.L.I. and D.H. conducted the data analysis, with the supervision of J.O. H.L.I. generated data visualizations. J.O. and H.L.I. wrote the paper with input from D.H.

## STAR Methods

### Resource Availability

#### Lead Contact

Further information and requests for resources and reagents should be directed to the Lead Contact, Julie Overbaugh.

#### Materials Availability

The data presented in this manuscript and research materials used in this study are available from the Lead Contact upon request.

#### Data and Code Availability

The custom RStudio code generated and used in this study to analyze and visualize data is available upon request. Any additional information required to reanalyze the data reported in this paper is also available upon request.

### Experimental model and subject details

#### Primary CD4^+^ T cells

Primary CD4^+^ T cell isolation, activation, single gene editing, and culturing procedures largely follow a previously published detailed protocol ^75^. For primary cell isolation, whole blood from healthy donors collected in EDTA tubes was obtained from BloodWorks Northwest. Peripheral blood mononuclear cells (PBMCs) were isolated by centrifugation over Ficoll-Paque Plus (Cytiva #17144002). CD4^+^ T cells were isolated using negative selection (STEMCELL Technologies #17952). CD4^+^ T cells were immediately cultured in complete RPMI media (RPMI 1640 supplemented with 10% FBS, 2 mM L-glutamine, 1X antibiotic-antimycotic that contains penicillin, streptomycin, and Amphotericin B) with 100 U/mL recombinant interleukin (IL)-2 (Roche #11147528001) and activated as described below or frozen in freezing media (90% FBS, 10% DMSO). For all procedures, cell counts were obtained using a hemocytometer.

#### Cell lines

HEK293T/17 (ATCC #CRL-11268) and TZM-bl cells (NIH HIV Reagent Program #ARP-8129) were cultured in DMEM complete media (DMEM supplemented with 10% FBS, 2 mM L-glutamine, 1X antibiotic-antimycotic that contains penicillin, streptomycin, and Amphotericin B).

#### Plasmids

pMD2.G (Addgene #12259) and psPAX2 (Addgene #12260) were gifts from Didier Trono. The TOPmCh*Δ*W vector ^49^ was modified to utilize it in HIV-CRISPR screens. First, the intact 3’UTR was reconstituted as described for the HIV-CRISPR vector ^22^. To do this, a gene fragment was synthesized (Integrated DNA Technologies (IDT)) and sub-cloned into the XcmI and PmeI sites using Gibson Assembly. Next, to allow for cloning in an sgRNA library and to incorporate an optimal tracrRNA sequence ^50^, a cassette was synthesized and assembled into the EcoRI and KpnI sites. In doing this, the spacer region between BsmBI sites used for cloning in of the sgRNA library was reduced from the 1885 bp of other commonly used CRISPR vectors to 355 bp. This modified version of the TOP vector is referred to as HIV-TOP opt mCh*Δ*W.

#### HIV-1 viruses

HIV Q23.BG505 ^37^ is a clade A, CCR5-tropic HIV chimeric virus derived from a full-length provirus isolated early in infection (Q23) ^36^ and bears the BG505.C2 envelope ^78^ of a well-studied clade A transmitted/founder virus. HIV LAI is a clade B, CXCR4-tropic HIV strain, as previously described ^38^. Q23.BG505 and LAI viral stocks were prepared by transfection as described below. The following viral stocks were obtained through the NIH HIV Reagent Program, Division of AIDS, NIAID, NIH: HIV Ba-L (ARP-510) contributed by Dr. Suzanne Gartner, Dr. Mikulas Popovic and Dr. Robert Gallo ^79^; HIV 93/MW/965 (ARP-2913) contributed by Dr. Paolo Miotti and the UNAIDS Network for HIV Isolation and Characterization; HIV 92/UG/029 (ARP-1650) contributed by UNAIDS Network for HIV Isolation and Characterization and the DAIDS, NIAID; HIV 92/UG/024 (ARP-1649) contributed by UNAIDS Network for HIV Isolation and Characterization.

### Method Details

#### Primary CD4^+^ T cell activation and culture

CD4^+^ T cells were used directly after isolation for the first set of single KO experiments (**Figure 2**) and for bulk RNA-seq experiments (**Figure 3**) and were thawed from frozen aliquots for all other experiments. After isolation or thawing, cells were resuspended at 2.5e10^6^ cells/mL in 48-well flat-bottom plates with complete RPMI media and 100 U/mL IL-2. Cells were activated with plate-bound anti-CD3 (Tonbo Biosciences #40-0038) at a concentration of 10 µg/mL and anti-CD28 (Tonbo Biosciences #40-0289) supplemented to the media at a concentration of 5 µg/mL.

#### RNP assembly for single- and multi-gene KOs

NTC RNPs utilized a single sgRNA from IDT, whereas RNPs for all other single gene KOs were assembled using a mixture of three sgRNAs from Synthego (Synthego Gene KO Kit v2). NTC RNPs were prepared in advance by preincubating 1 µL of 160 nM CRISPR-RNA (crRNA, IDT) with 1 µL of 160 nM tracrRNA (IDT) at 37C for 30 min. The crRNA/tracrRNA complex was then incubated for 15 min at 37C with 2 µL of 40 nM Cas9- NLS protein (UC Berkeley QB3 MacroLab) for a final molar ratio of 4:1 RNA:Cas9. Complexes were frozen at - 80C until ready for use. For all other single gene KOs, RNPs were assembled on the day of nucleofection by combining 6 µL of 30 pmol/µL sgRNAs (Synthego), 18 µL of P3 Primary Cell Nucleofector solution (Lonza #V4SP-3096), and 1 µL of 20 µM Cas9-NLS protein (UC Berkeley QB3 MacroLab). Complexes were gently mixed, incubated for 10 min at room temperature, and then stored on ice until ready for same-day use.

For multi-gene KOs, RNPs for each gene were prepared separately and respective to the number of genes being targeted. For each gene, 1.5 µL of 60 pmol/µL sgRNAs (Synthego) was combined with 1 µL of 20 µM Cas9-NLS protein (UC Berkeley QB3 MacroLab) and the following volume of P3 Primary Cell Nucleofector solution: 8.75 µL for two-gene KOs, 5.67 µL for three-gene KOs, and 3.2 µL for five-gene KOs. Complexes were gently mixed, incubated for 10 min at room temperature, and then stored on ice until ready for same-day use. Immediately prior to nucleofection, RNPs for different genes were combined.

#### RNP nucleofection of CD4^+^ T cells

Three days after activation, 1e6 CD4^+^ T cells per nucleofection were pelleted at 100xg for 10 min and washed once with PBS. For nucleofections with NTC RNPs, cells were resuspended in 20 µL of P3 Primary Cell Nucleofector solution (Lonza #V4SP-3096) and mixed gently with 4 µL of the complexes described above. For RNPs with Synthego guides (both single and multi-gene KO), cells were gently resuspended with 25 µL of the complexes described above. Cells and complexes were incubated for 5 min at room temperature before nucleofection in a 16-well Nucleocuvette (Lonza #V4SP-3096) in an Amaxa Nucleofector (Lonza) using pulse code EH-115. Cells were supplemented with 80 µL of RPMI complete media with 125 U/mL IL-2 and allowed to recover in the cuvette for 30 min to 2 h at 37C. Cells were then brought to 2.5e6 cells/mL in RPMI complete media with 100 U/mL IL-2 and plated in 96-well flat-bottom plates at a 1:1 ratio with activation beads (Miltenyi Biotec #130-091-441). Two days later, 200 µL of RPMI complete media with 100 U/mL IL-2 was added to each nucleofection. At four days post nucleofection, fresh media was added to each nucleofection by pelleting cells at 300xg for 10 min, removing 250 µL of supernatant (roughly half of the total culture volume), and adding RPMI complete media with 100 U/mL IL-2 to bring cells to 1e6 cells/mL. Starting the following day, cells were counted and resuspended to 1e6 cells/mL in RPMI complete media with 100 U/mL IL-2 every other day. As needed, cells were transferred from 96-well plates (working volume: 100-300 µL/well) to 48-well plates (400-600 µL/well) or 24-well plates (1-2 mL/well).

#### HIV-1 and lentivirus production and concentration

HIV and lentiviral stocks were prepared by transfecting HEK293T/17 cells (ATCC #CRL-11268) in a six-well format. Twenty hours prior to transfection, 5e5 cells/well were plated in 2 mL of DMEM complete. On the day of transfection, serum-free DMEM media, transfection reagents, and plasmids were equilibrated to room temperature. For each HIV transfection well, 1 µg of proviral DNA and 200 µL serum-free DMEM were combined, prior to the addition of 18 µL of FuGENE (Promega #E2692). For each lentiviral transfection well, 667 ng of transfer plasmid, 500 ng of psPAX2 packing vector (Addgene #12260), 333 ng of the pMD2.G VSV-G expression vector (Addgene #12259), and 200 µL serum-free DMEM were combined, prior to the addition of 4.5 µL TransIT- LT1 transfection reagent (Mirus Bio #MIR-2305). Transfection complexes were gently mixed, incubated for 15-30 minutes at room temperature, and added to each well in a drop-wise manner. At 20 h post-transfection, media was replaced with 1.5 mL of DMEM complete per well. Supernatants were harvested two days post-transfection, removed of debris using a 0.2 µM filter (Millipore Sigma #SCGP00525), and concentrated as described below.

For HIV transfections, supernatants were passed through Amicon 100 KDa concentrators (Millipore Sigma #UFC910024) until concentrated ∼100-fold. Concentrated virus was aliquoted and stored at -80C. HIV stocks were titered on TZM-bl cells (NIH HIV Reagent Program #ARP-8129). Infections were carried out in the presence of 10 µg/mL DEAE-dextran and viral titer was determined by staining fixed cells for β-galactosidase activity at 48 h post-infection and counting blue, infected cells.

For lentiviral transfections, ice-cold PBS was added to supernatants to bring the volume to 35 mL. Diluted supernatants were concentrated over 2 mL of cold 20% sucrose by ultra-centrifugation at 23,000 rpm for 1 h at 4C. After pouring the supernatant off, the sucrose pellet was resuspended in RPMI complete media, such that the virus was concentrated ∼100-fold. Pellets were resuspended by periodically vortexing at a low intensity and were left at 4C overnight before aliquoting and freezing at -80C. Lentiviral stocks were titered by copy number as described below.

#### IFN treatment and HIV infection of CD4^+^ T cells

Nucleofected CD4^+^ T cells were treated with 1000 U/mL IFN-*β*1a (PBL Assay Science #11410) 24 h prior to HIV infection. For the first set of single KO experiments (**Figure 2**) and bulk RNA-seq experiments (**Figure 3**), cells were IFN-treated six days post nucleofection. Otherwise, cells were IFN-treated nine days post nucleofection to allow for additional recovery. For targeted KO experiments, half of each nucleofection was IFN-treated, whereas all cells were IFN-treated for CRISPR screens. For ISG KO spreading infections, cells were resuspended to 1e6 cells/mL in RPMI complete media with 100 U/mL IL-2 and 100 µL was added in duplicate for each virus to 96- well flat-bottom plates. Cells were infected with 10 µL of virus at an MOI of 0.02 in the presence of 8 µg/mL polybrene (Millipore Sigma #TR-1003-G) by spinoculation at 1100xg for 90 min at 30C. After infection, cells were transferred to 96-well V-bottom plates, pelleted at 300xg for 10 min, and resuspended in 100 µL of fresh RPMI complete with 100 U/mL IL-2 that contained 1000 U/mL IFN as necessary. Cells were then transferred back to the 96-well flat-bottom infection plates and were incubated at 37C. Viral supernatants were harvested 2, 4, 6, and 8 days post-infection (dpi), stored at -80C, and replaced with RPMI complete media with 100 U/mL IL-2 containing 1000 U/mL IFN as needed.

#### Reverse transcriptase (RT) activity qPCR assay

HIV infection levels were assessed by measuring RT activity in viral supernatants using the previously described RT activity qPCR assay ^80^. Upon thawing, 5 µL of viral supernatant was incubated with 5 µL lysis buffer (0.25% Triton X-100, 50 mM KCl, 100 mM Tris HCl, 40% glycerol, 0.8 U/µL RNAse Inhibitor (ThermoFisher #EO0381)) for 10 min at room temperature. This mixture was then diluted with 90 µL of ultrapure water and 9.6 µL was transferred to a 384-well MicroAmp Optical plate (Applied Biosystems #4309849) as input for the assay. Per well, 10.4 µL of the following reaction mix was used: 10 µL ROX SYBR 2X MasterMix (Eurogentec #UF-RSMT- B0701), 0.1 µL 10-fold diluted RNase Inhibitor (ThermoFisher #EO0381), 0.1 µL of MS2 RNA (Roche #10165948001), and 0.1 µL of each 100 µM primer (MS2-F: TCCTGCTCAACTTCCTGTCGAG, MS2-R: CACAGGTCAAACCTCCTAGGAATG). Plates were read on an Applied Biosystems QuantStudio 7 Real Time PCR machine.

To quantify the RT activity (mU/mL) of viral supernatants, the RT units of a concentrated stock of HIV Q23.BG505 virus were determined multiple times using a standard curve of purified HIV RT enzyme (Worthington Biochemical Corp. #LS05003). Aliquots of this titered stock of Q23.BG505 were then used as the quantitative standard curve in all assays.

#### Genomic editing analysis

On the day of HIV infection, 2.5e5 nucleofected cells were harvested by pelleting at 300xg for 10 min in PBS. Cell pellets were either frozen at -80C or immediately used for genomic DNA extraction (QIAGEN #51104). Edited loci were amplified from genomic DNA from edited and control cells using 10 µM primers specific to each targeted locus (**Table S5**), Herculase II Fusion DNA Polymerase (Agilent #600679), and the thermocycler program detailed in **Table S5**. PCR amplicons were purified (QIAGEN #28104) and Sanger sequenced (Fred Hutch Shared Resources Genomics Core). Chromatograms were analyzed by ICE analysis tool v3 (Synthego) to determine the rate of editing predicted to cause KO at each genomic locus in the cell population.

#### Bulk RNA-seq of IFN-treated CD4^+^ T cells

CD4^+^ T cells were isolated and immediately activated on the same day, as described above. Three days later, cells were brought to 2.5e6 cells/mL in RPMI complete media with 100 U/mL IL-2 and plated in 96-well flat-bottom plates at a 1:1 ratio with activation beads (Miltenyi Biotec #130-091-441). Cells were maintained thereafter as usual. Six days after bead activation, CD4^+^ T cells were treated with 1000 U/mL IFN-*β*1a (PBL Assay Science #11410) as described above. Cells were collected prior to IFN treatment and 3, 6, 12, and 24 h thereafter, depending on the donor (**Figure 3A**). During collection, cells were washed with PBS, pelleted, lysed in 350 µL Buffer RLT Plus (QIAGEN #74134) supplemented with 1% *β*-mercaptoethanol, and stored at -80C. This procedure was repeated for five independent donors.

For RNA extractions, thawed cell lysates were homogenized using a QIAshredder column (QIAGEN #79654) and genomic DNA was removed on a gDNA Eliminator column. Total RNA was then extracted using the RNeasy Plus Mini Kit (QIAGEN #74134) and eluted in 30 µL RNase-free water. The quality and concentration of RNA preps was assessed by Agilent High Sensitivity RNA ScreenTape (Fred Hutch Shared Resources Genomics Core). After samples were found to have high RNA Integrity scores (RIN ≥ 9.4), they were sequenced on an Illumina HiSeq 2500 (Fred Hutch Shared Resources Genomics Core) with a sequencing depth of roughly 20 million reads per sample.

#### CD4-ISG sgRNA library construction

An oligo pool containing the CD4-ISG sgRNA library was synthesized (Twist Biosciences) and cloned into HIV- TOP opt mCh*Δ*W, as previously described in detail ^22^. The library pool was amplified (Array-F primer: TAACTTGAAAGTATTTCGATTTCTTGGCTTTATATATCTTGTGGAAAGGACGAAACACCG, optTracr-Array-R primer: GTTGATAACGGACTAGCCTTATTTCAACTTGCTATGCTGTTTCCAGCATAGCTCTGAAAC), gel-purified (QIAGEN #28704), and cloned into the HIV-TOP opt mCh*Δ*W following digestion with BsmBI (NEB #R0739L) using Gibson Assembly (NEB #E2611S) with a 5:1 molar ratio of insert to vector. Gibson reactions were transformed into Endura electrocompetent cells (Lucigen #60242-2) at a large scale, such that there was 230-fold coverage of the CD4-ISG library. Transformed bacteria were scraped from plates and plasmid DNA was isolated using the Endotoxin-Free Nucleobond Plasmid Midiprep Kit (Takara Bio #740422.10). The CD4-ISG library was sequenced and contains 4,574 of the 4,759 guides (99.9%) initially included in library design.

#### Titration of lentivirus stocks by copies per cell

CD4^+^ T cells were isolated, thawed, and activated as described above but were plated in a 96-well plate. The next day (20-24 h), cells were transduced with a dilution series of the concentrated CD4-ISG library lentivirus. Each well was transduced with 10 µL of viral inoculum supplemented with protamine sulfate (Millipore Sigma #P3369) for a final concentration of 8 µg/mL. Cells were spinoculated at 1100xg for 90 min at 30C and maintained at 37C afterwards. Four days after transduction, roughly 1e5 cells were collected, pelleted in PBS, and subjected to genomic DNA isolation (QIAGEN #51104). Genomic DNA was digested with BglI (NEB #R0143L) and diluted 1:3 for droplet digital PCR (ddPCR) amplification to measure vector copies per cell. Per well of a 96-well ddPCR plate (Bio-Rad, #12001925), 3 µL of diluted template DNA was used as input, along with 1.8 µL of each 10 µM vector-specific primer (ddPCR-cPPT-F: GTACAGTGCAGGGGAAAG, ddPCR-U6-R: ATGGGAAATAGGCCCTCG), 0.5 µL FAM probe (6-FAM/ZEN - AGACATAATAGCAACAGACATACAAAC - IBFQ), 1 µL RPP30 ddPCR copy number variation HEX assay mix (Bio-Rad, #100-31243), and 10 µL SuperMix with no dUTPs (Bio-Rad #1863024). The volume per well was brought to 20 µL with water, followed by droplet generation, PCR amplification, and reading of the plate on an Applied Biosystems QuantStudio 7 Real Time PCR machine. Vector copies per cell were calculated by dividing vector copies/µL by RPP30 copies/µL, and then dividing this value by two to correct for two RRP30 copies per cell. Titrations were performed in CD4^+^ T cells from two unique donors during independent experiments. Cumulative results determined that 0.06 µL of CD4-ISG library lentivirus per 1e6 cells yielded an average of 2.7 vector copies per cell.

#### CD4-ISG HIV-CRISPR screening

CD4^+^ T cells were isolated, thawed, and activated as described above. The next day (20-24 h), roughly 20e6 cells were transduced with the CD4-ISG library lentivirus, aiming to deliver 3 vector copies per cell as described in titration experiments above. Each well was transduced with 10 µL of diluted viral inoculum supplemented with protamine sulfate (Millipore Sigma #P3369) for a final concentration of 8 µg/mL. Cells were spinoculated at 1100xg for 90 min at 30C and maintained at 37C afterwards. Two days later, Cas9 was delivered to cells by nucleofection with NTC RNPs, as described above. This approach utilizes a guide-swap strategy ^52^ that replaces the nucleofected NTC guide with the library guides delivered by lentiviral transduction. Cells were maintained thereafter as usual. Nine days after nucleofection, all library cells were treated with 1000 U/mL IFN-*β*1a (PBL Assay Science #11410). On the following day, HIV infections were established.

For infections, 2.5e5 CD4^+^ T cells in 100 µL of RPMI complete media with 100 U/mL IL-2 and 1000 U/mL IFN were added to wells of a 96-well flat-bottom plate for each virus and MOI. In total, 10e6 IFN-treated CD4^+^ T cells were used for each CRISPR screen to maintain high library coverage. Cells were infected with 10 µL of virus at an MOI of 0.1 and 1 in the presence of 8 µg/mL polybrene (Millipore Sigma #TR-1003-G) by spinoculation at 1100xg for 90 min at 30C. After infection, 150 µL of RPMI complete media with 100 U/mL IL-2 and 1000 U/mL IFN was added to each well to bring cells to 1e6 cells/mL and plates were incubated at 37C. Three days post infection, all cells were split 1:2 with fresh RPMI complete media with 100 U/mL IL-2 and 1000 U/mL IFN. For high MOI screens (HIV MOI=1), viral supernatants were harvested 3 and 5 dpi and cells were collected 5 dpi. For low MOI screens (HIV MOI=0.1), viral supernatants were harvested 4 and 7 dpi and cells were collected 7 dpi. All viral supernatants were filtered to remove cell debris (Millipore Sigma #SCGP00525) and stored at 4C. Within four days of collection, viral supernatants were concentrated over a sucrose gradient as described above, resuspended in 100 µL PBS, and stored at 4C. Viruses harvested on different days were kept separate throughout this process. Cells were washed in PBS, pelleted, and stored at -80C.

CRISPR screen samples were prepared for Illumina sequencing as previously described ^22^. Genomic DNA was extracted from thawed cell pellets using the QIAamp DNA Blood Midi Kit (QIAGEN #51185) and digested with BglI (NEB #R0143L). Viral RNA was isolated from viral pellets one day after concentration (QIAGEN #52904). RNA extracts were then treated with DNase I (Roche #04716728001) and reverse transcribed to generate cDNA with Superscript III (Invitrogen #18080044) and a primer specific to the HIV-TOP opt mCh*Δ*W vector (SeqMSCV- R1-R: CTTGCTAAACCTACAGGTGG). Guide sequences were amplified from 20 µg of prepared genomic DNA (∼400-fold coverage) and viral cDNA using Herculase II Fusion DNA Polymerase (Agilent #600679) and 10 µM primers specific to the HIV-TOP opt mCh*Δ*W vector (PLC-seq-R1-F: GAGGGCCTATTTCCCATGATTCCTTCA, SeqMSCV-R1-R: CTTGCTAAACCTACAGGTGG). Amplicons were purified and concentrated (QIAGEN #28104). A second-round PCR was performed to add identifier sequences to each sample using indexing primers (**Table S5**) and the PLC-Seq-R2-RS primer (CAAGCAGAAGACGGCATACGAGATGTGACTGGAGTTCAGACGTGTGCTCTTCCGATCTTGCCACTTTTTCAAGTTGATAACGGACT). PCR products were gel isolated (QIAGEN #28704) and quantified using the Quant-iT PicoGreen dsDNA Assay Kit (Invitrogen #P7589). Samples were pooled at an equimolar ratio to generate a 2 nM pool. Pooled libraries were sequenced on an Illumina NextSeq 2000 with ≥500-fold coverage (Fred Hutch Shared Resources Genomics Shared Resource).

#### Western blotting

Cells collected on the day of HIV infection were lysed in RIPA buffer (50 mM Tris pH 8.0, 0.1% SDS, 1% Triton-X, 150 mM NaCl, 1% deoxycholic acid, 2 mM PMSF). Cell extracts were prepared in 4X LDS Buffer (Invitrogen #NP007) and 2X MES SDS Running Buffer (Invitrogen #NP002). After boiling at 100C for 5 min, 20 µg of the prepared samples were loaded into a NuPAGE 4-12% Bis-Tris Gel (Invitrogen #NP0321) and resolved. Following transfer onto a nitrocellulose membrane (Invitrogen #LC2001), blots were blocked in blocking buffer (PBS, 0.1% Tween, 5% milk) for 1 h, incubated with the MX2 primary antibody (Santa Cruz Biotech #sc-271527; 1:100) overnight, probed with secondary antibody (Cytiva #NA931, 1:1000) for 1 h, and visualized by chemiluminescence (Cytiva #RPN2232) on a Bio-Rad ChemiDoc Touch. Membranes were washed in PBST (PBS, 0.1% Tween) in between each step. After imaging the MX2 stain, membranes were stripped (ThemoFisher #46430), washed, probed with GAPDH:HRP (Bio-Rad #MCA4739P, 1:1000) for 1 h, and imaged with chemiluminescence.

#### HIV CA sequence alignments

CA sequences were obtained via http://www.hiv.lanl.gov/. Global alignments were performed using Geneious Prime version 2022.2 (RRID:SCR_010519).

### Quantification and Statistical Analysis

#### RT activity qPCR assay analysis

The RT activity (mU/mL) of experimental wells was interpolated from a standard curve (concentrated Q23.BG505 stock, as described above) using the lm function in the R stats package. RT values from duplicate experimental wells were then averaged prior to analysis. To compare RT activity across donors for KO experiments, the following were determined for each KO, virus, and donor using day 6 RT activity: Relative IFN Sensitivity (no IFN RT / IFN RT), the fold increase in HIV Levels with IFN due to ISG KO as compared to the negative control (ISG KO / negative control), and the fold decrease in Relative IFN Sensitivity due to ISG KO as compared to the negative control (negative control / ISG KO). NTC-delivered cells were used as the negative control for the first set of single KOs (**Figure 2**). For all other KO experiments (**Figures 5 and 6**), we inactivated the B cell marker CD19 as the negative control because nucleofection with CD19 RNPs generates double-stranded DNA breaks, making it an even more robust negative control, and does not impact HIV replication, which we verified (**Figure S6**). Standard curve interpolation and statistical comparisons were performed in RStudio.

#### Bulk RNA-seq analysis and CD4-ISG library design

Illumina sequencing reads (SRA accession number: PRJNA921704) were aligned to the GRCh38 reference genome to generate raw counts (Fred Hutch Shared Resources Bioinformatics Core). Counts per million (CPM) were then calculated using the cpm function in the Bioconductor edgeR package ^81^ (**Table S1**). For principal component analysis (PCA) of all 17 samples, data from genes with sufficient expression (CPM>1 in at least 8 of the samples) were corrected for donor variation with the combat_seq function ^82^ in the Bioconductor sva package prior to PCA generation.

Prior to calculating IFN induction, raw counts equal to 0 were adjusted to 1 and CPMs were re-generated. With roughly 20 million reads per sample, cases originally with zero counts ultimately had ∼0.05 CPM, which allowed for fold change calculations but should not skew downstream analyses. After CPM generation, long intergenic non-coding RNAs genes (GeneName contains “LINC”) were removed as were mitochondrial genes (Chr = MT). ISG designation was then performed for each time point independently. For data from 6, 12, and 24 hours after IFN treatment, CPMs from IFN-treated samples from a given time point and their donor-matched untreated controls were aggregated and genes with low expression were filtered out (required CPM>1 in half of the samples). IFN fold induction was calculated from CPMs for each treated/untreated donor pair (**Table S1**). Genes with a fold change (FC) ≥2 in the majority of donors for that time point were considered ISGs. This yielded 449, 261, and 167 ISGs for the 6, 12, and 24 h post IFN time points, respectively. For data from 3 hours after IFN treatment, we implemented a more stringent strategy for ISG designation because we only had data from one donor for this time point. IFN fold induction was calculated for the 3 h treated/untreated donor pair for genes that passed the CPM cutoff for any of the prior time point analyses. Genes with a FC≥4 were considered ISGs, yielding 261 ISGs. These analyses identified 573 genes in total (**Figure 3D**). Finally, to capture genes with high IFN stimulation in any donor or time point, we also included genes with FC≥5 in any sample, which added 27 genes for a total of 600 ISGs.

To design the CD4-ISG library, we generated four guides with two different tools, CHOPCHOP ^83^ and GUIDES ^84^, for a total of eight guides per gene. Of the 600 ISGs identified via bulk RNA-seq, we were unable to design guides for 107 due to transcripts encoding pseudogenes, RNA genes, or read-through regions. To complete the CD4-ISG sgRNA library, we added 21 control genes, 2 genes from the first set of single KOs, 38 top-scoring hits from preliminary CRISPR screens in CD4^+^ T cells with the PIKA ISG library ^22^ (data not shown), and 200 NTC guides for a total of 555 genes represented across 4,579 guides (**Table S2**).

#### CD4-ISG HIV-CRISPR screen analysis

Illumina sequencing reads (SRA accession number: PRJNA919035) were analyzed by MAGeCK-Flute ^53^ to generate guide counts and gene-level enrichment data (**Table S3**; Fred Hutch Center for Data Visualization). As opposed to using all 200 NTC guides to inform data for one NTC “gene”, NTC guide sequences were iteratively binned to create a number of NTC “genes” (synthetic NTCs, synNTCs) with eight guides per gene to match the number of genes in the CD4-ISG library (554) (**Table S2**). This approach captures variation across NTC guides when analyzing gene-level data more accurately than aggregating all 200 guides into one NTC gene-level data point. Results from viral supernatants collected 3 dpi for high MOI screens and 4 dpi for low MOI screens were used for the analyses presented in this report, as these collection days had higher donor correlation coefficients compared to the later collection time points (5 and 7 dpi, respectively by MOI; data not shown). Pearson R correlations between MAGeCK scores from different donors were determined using the stat_cor function in the ggpubr package. Screen background was calculated as the average of the synNTC MAGeCK scores + 3 standard deviations.

## KEY RESOURCE TABLE

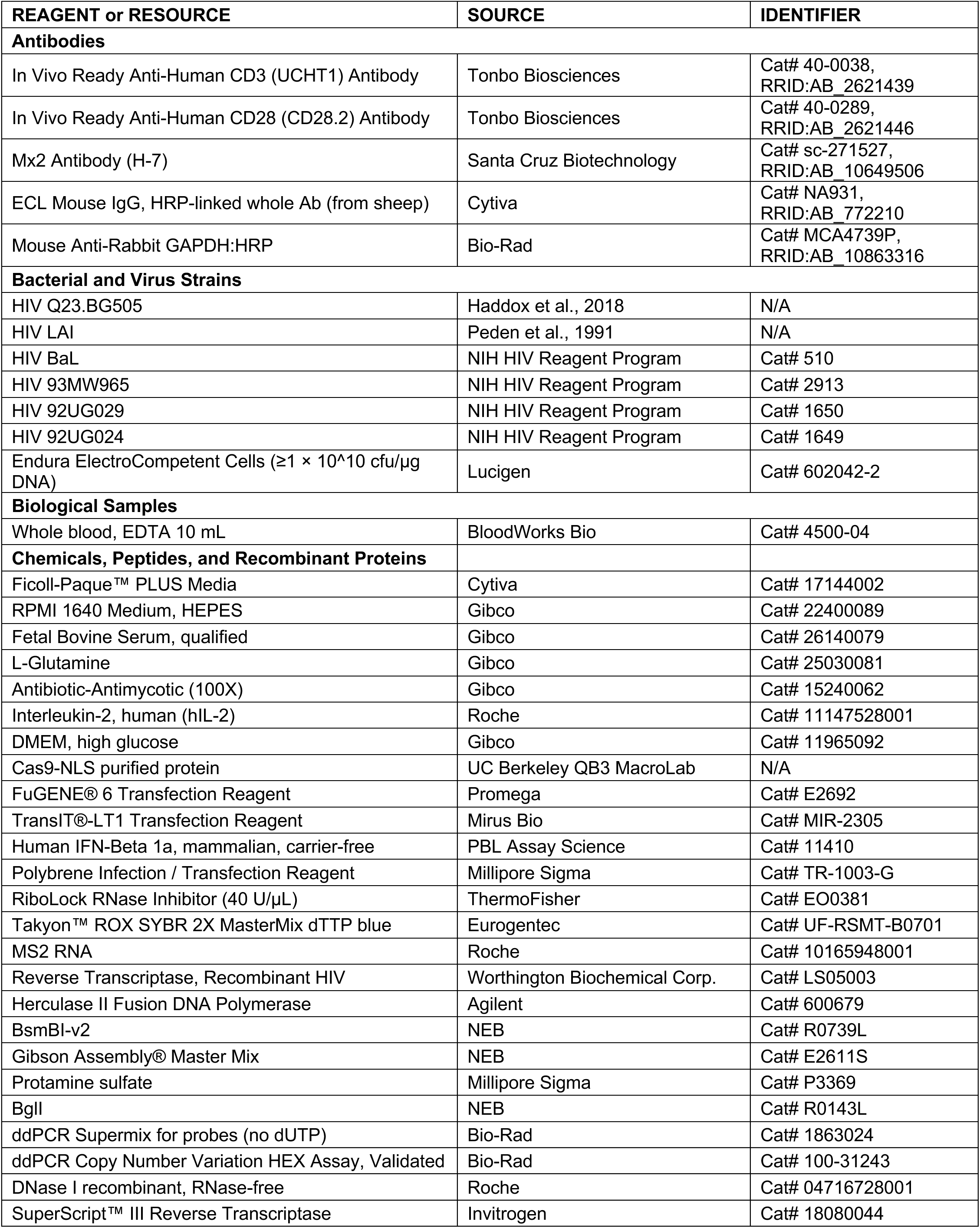
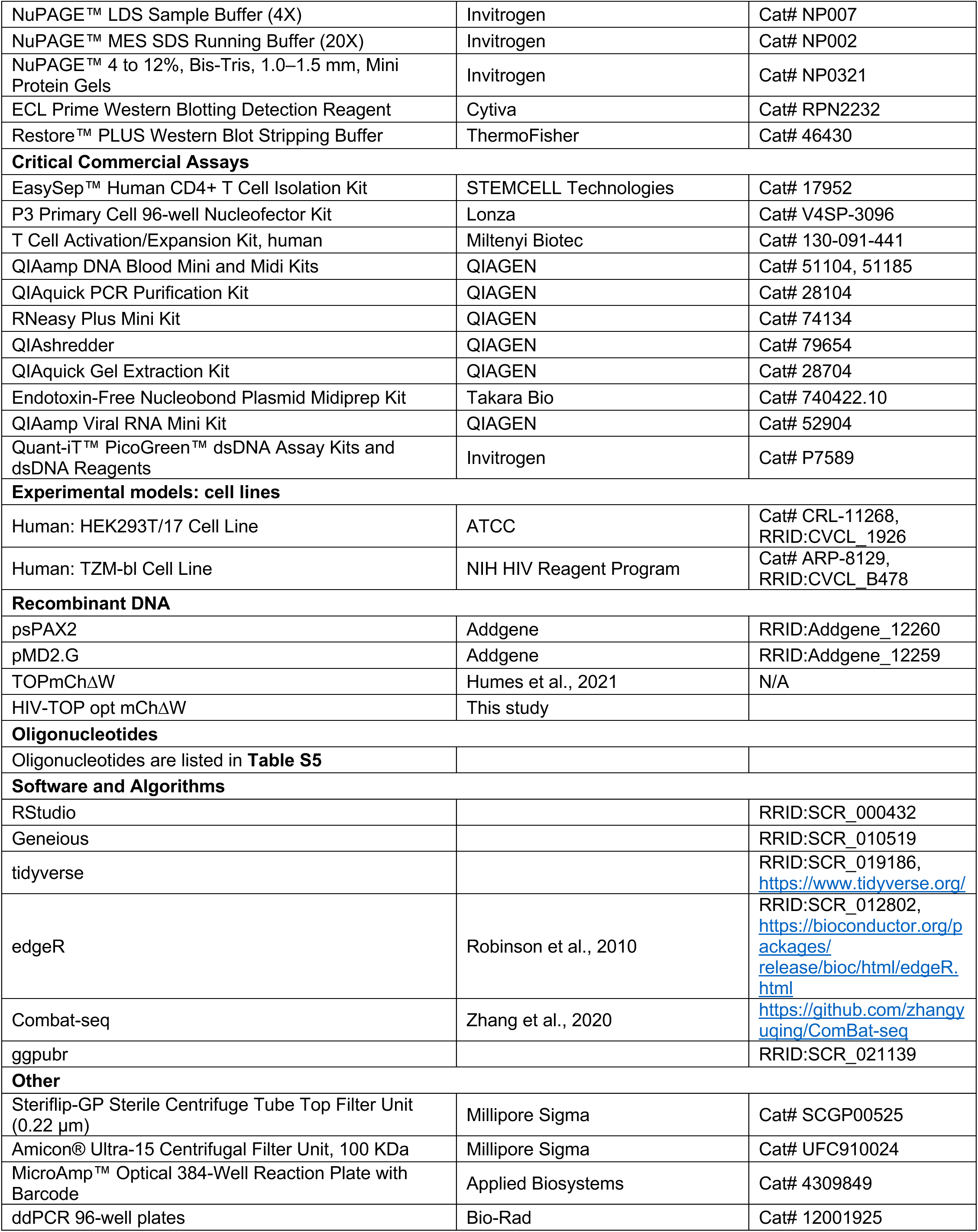

