## Supplementary material for "Several cell-intrinsic effectors drive type I interferon-mediated restriction of HIV-1 in primary CD4^+^ T cells": Figure S1

**Figure S1, related to Figure 1.**

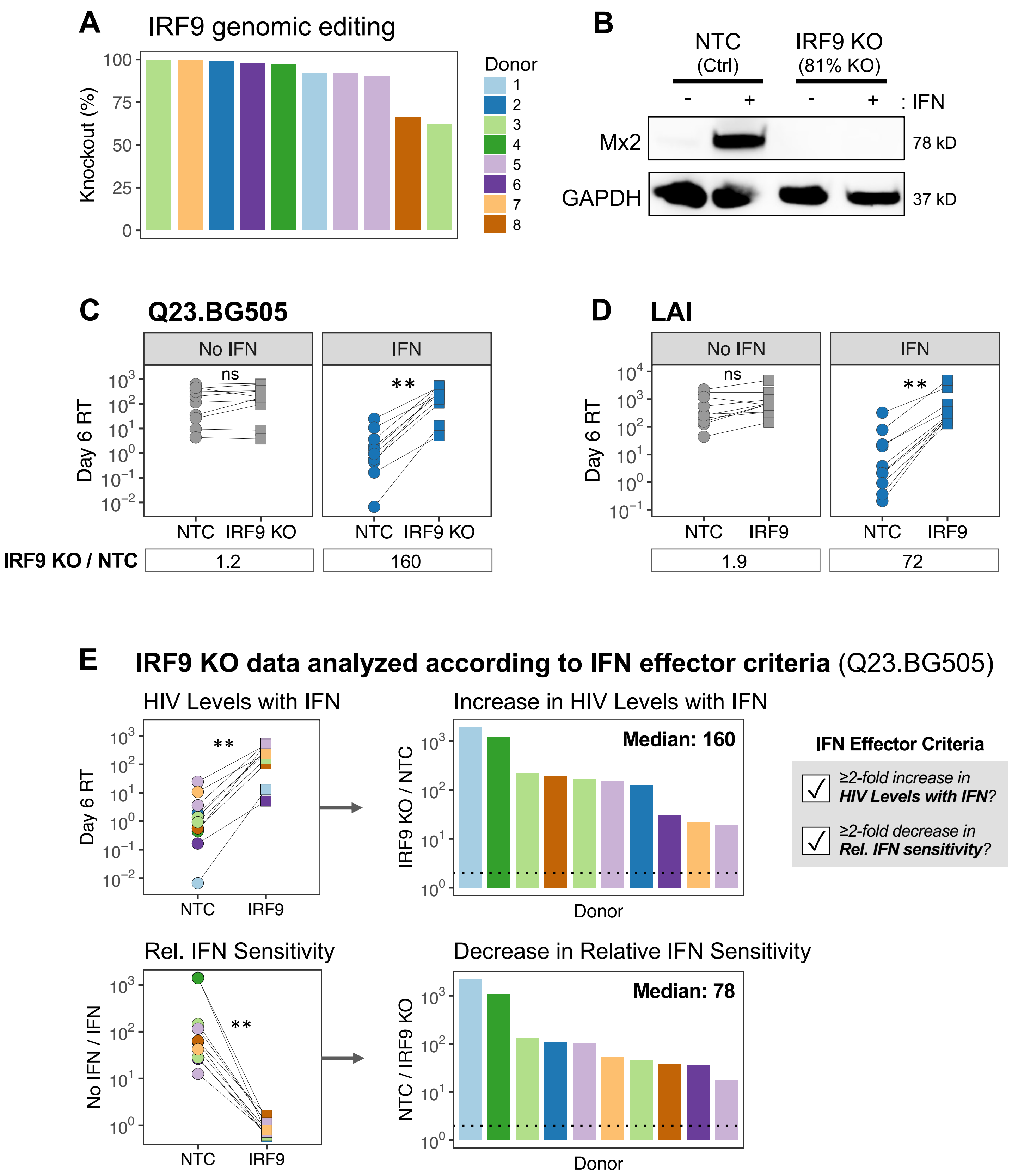

**Figure S1, related to Figure 1. IRF9 editing ablates the antiviral ISG response in primary CD4$^+$ T cells.** (**A**) IRF9 RNP-delivered CD4$^+$ T cells were collected on the day of infection and assessed for genomic editing by Sanger sequencing. Synthego ICE analysis was applied to sequencing results to determine percent knockout. (**B**) Western blot with lysates from NTC and IRF9-edited CD4$^+$ T cells from one donor to evaluate protein levels of the highly induced ISG MX2. (**C, D**) Paired data from Figure 1B and 1C, reformatted to compare infection results between NTC and IRF9-edited cells with and without IFN treatment. (**E**) HIV infection levels according to the metrics used to evaluate whether ISGs are IFN effectors. Left, paired data from panel B reformatted to compare "HIV Levels with IFN" and "Relative IFN Sensitivity" between NTC and IRF9-edited cells. Right, fold impact of IRF9 KO on these metrics, with bars depicting independent experiments. The median fold difference is denoted in the top right corner. The dotted line indicates a fold difference of two. **$p<0.002$, paired Wilcoxon rank test. Non-significant (ns) indicates $p>0.05$.
