## Supplementary material for "Several cell-intrinsic effectors drive type I interferon-mediated restriction of HIV-1 in primary CD4^+^ T cells": Figure S2

**Figure S2, related to Figure 2.**

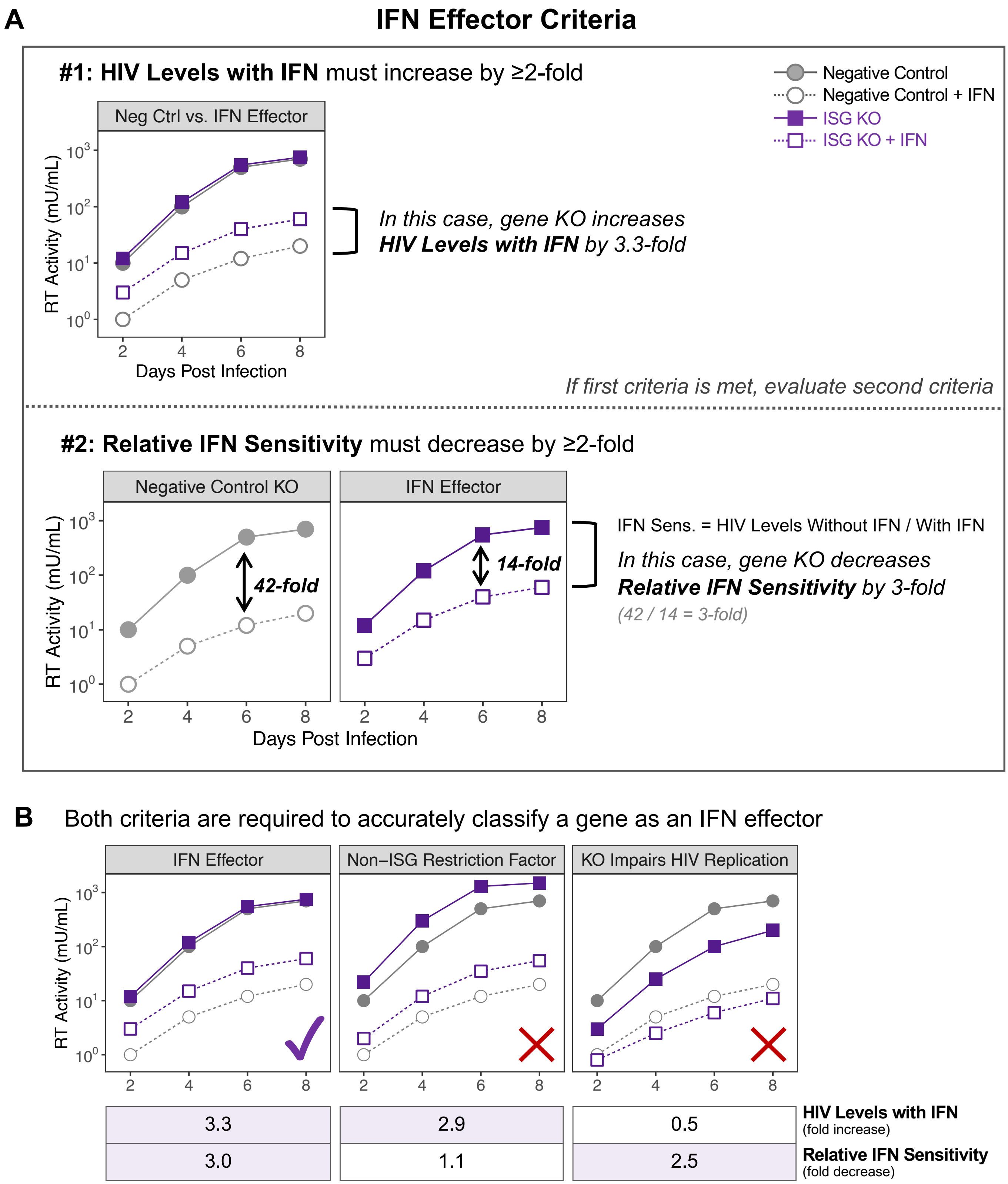


**Figure S2, related to Figure 2. Schematic of IFN effector criteria.**

**(A)** The depicted hypothetical HIV infection curves serve as examples to illustrate how ISG KO data was compared to negative control results to categorize genes as IFN effectors. RT activity measured on 6 dpi are used to calculate fold changes compared to an intra-assay negative control. (**B**) Example of an IFN effector and two types of genes that might impact these criteria but would not be classified as IFN effectors.
