## Supplementary material for "Several cell-intrinsic effectors drive type I interferon-mediated restriction of HIV-1 in primary CD4^+^ T cells": Figure S3

### Figure S3, related to Figure 2

**A** HIV Levels with IFN

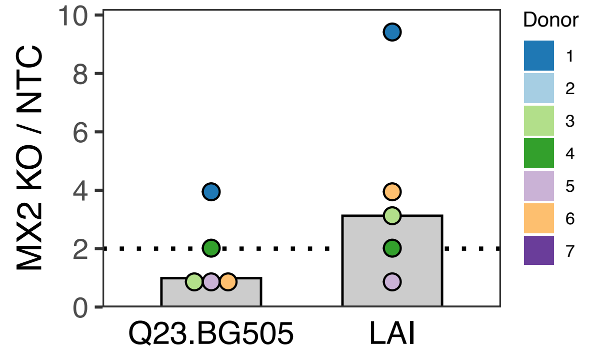


**B** Viruses tested for MX2 sensitivity

| Virus | MX2 Sensitive | Clade | Tropism | Primary Isolate | Ref |
|---|---|---|---|---|---|
| **Q23-17** | **No** | **A** | R5 | Yes | This study |
| CH040.c | **No** | B | R5 | Yes | 35 |
| RHPA.c | **No** | B | R5 | Yes | 35 |
| LAI/IIIB | Yes | B | X4 | No | This study, 21, 22 |
| NL4-3 | Yes | B | X4 | No | 21, 26, 30, 31, 35, 45, 46, 47, 48 |
| CH058.c | Yes | B | R5 | Yes | 35 |
| CH077.t | Yes | B | R5 | Yes | 21, 35 |
| CH106.c | Yes | B | R5 | Yes | 21, 35 |
| REJO.c | Yes | B | R5 | Yes | 21, 35 |
| YU-2 | Yes | B | R5 | No | 21 |
| WITO.c | Yes | B | R5 | Yes | 35 |

**C** HIV CA sequence alignment

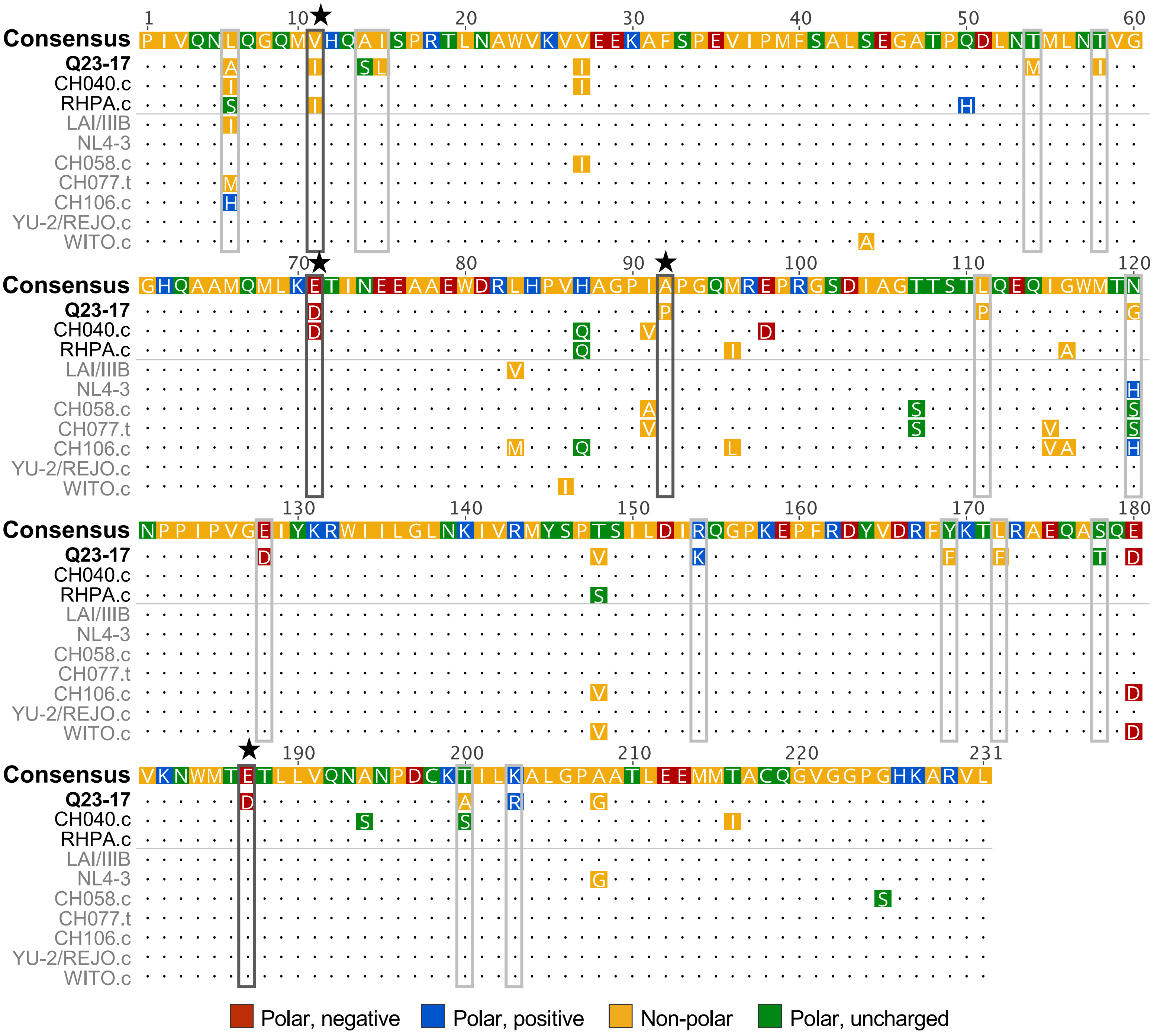

**Figure S3, related to Figure 2** (continued)

**D** HIV CA amino acid conservation

| *aa position* | 11 | 71 | 92 | 187 |
|---|---|---|---|---|
| **Q23-17** | **I** | **D** | **P** | **D** |
| **Group M** | V | **D** | **P** | E |
| **Clade A** | **I** | **D** | **P** | E |
| **Clade B** | V | E | A | E |
| **Clade C** | V | **D** | A | **D** |
| **Clade D** | V | E | A | E |
| **CRF01_AE** | V | E | **P** | E |
| **CRF02_AG** | V | **D** | **P** | E |
| **CRF07_BC** | V | **D** | A | **D** |

**Figure S3, related to Figure 2. HIV Capsid (CA) amino acid residues differ between Q23-17 and MX2-sensitive viruses.** (**A**) Increase in HIV Levels with IFN as a result of MX2 KO compared to NTC (n=5 donors, data also in Figure 2). Donor colors correspond to those in Figure 2. (**B**) HIV strains assessed for MX2 sensitivity in the current report and previous studies. (**C**) Alignment of HIV CA protein sequences from viruses in panel B. HIV strains are colored according to MX2 sensitivity (black: MX2-resistant, gray: MX2-sensitive). Amino acids are colored based on polarity. Locations in which the Q23-17 residue differs from all MX2-sensitive strains are boxed. Black, starred boxes indicate the four residues most likely to contribute to Q23-17's MX2 resistance. (**D**) Conservation of amino acids (aa) at sites likely to drive Q23.BG505's resistance to MX2 restriction. HIV group and clade consensus sequences are based on Troyano-Hernáez et al., 2022.
