## Supplementary material for "Several cell-intrinsic effectors drive type I interferon-mediated restriction of HIV-1 in primary CD4^+^ T cells": Figure S4

**Figure S4, related to Figure 4.**

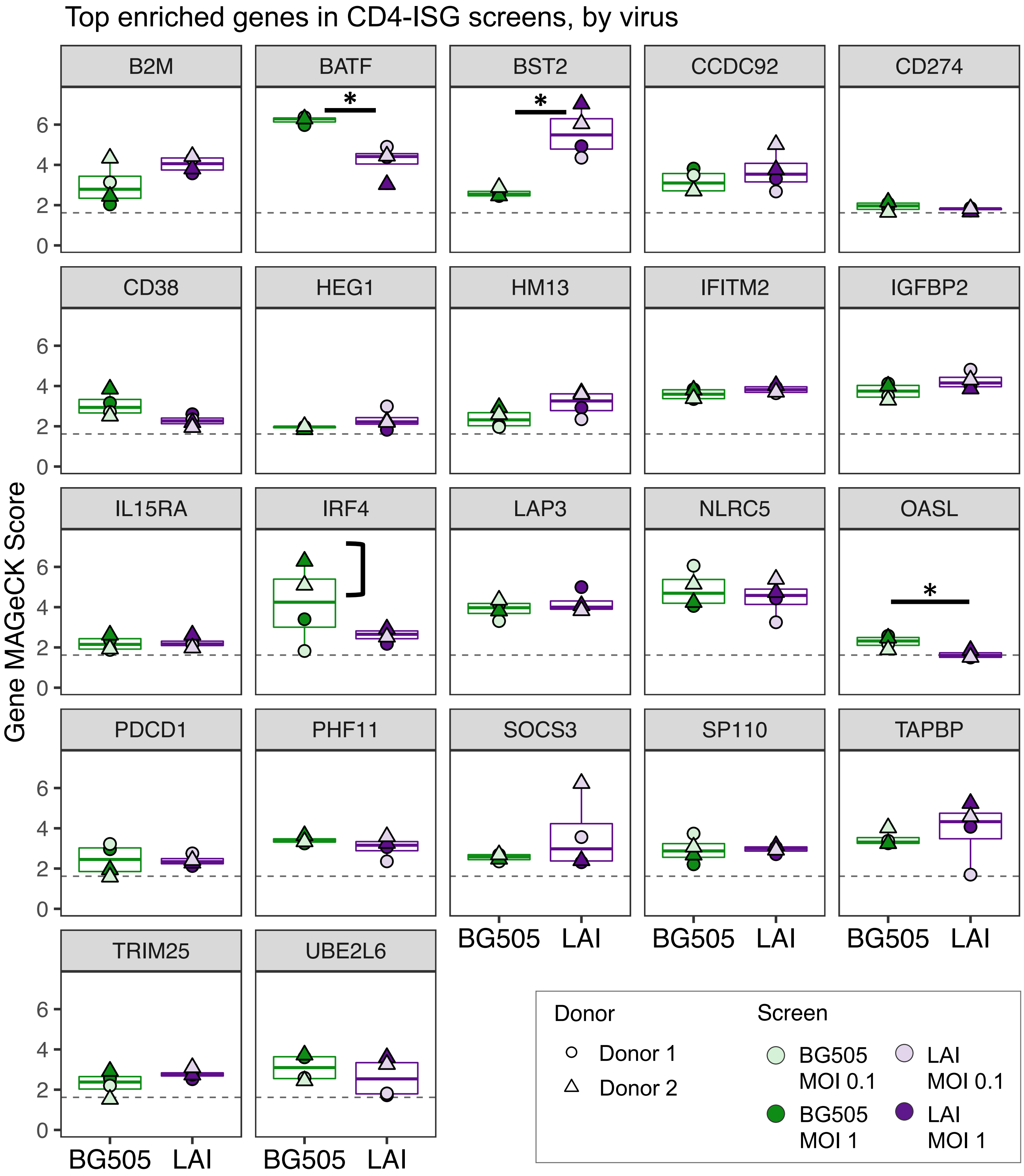


**Figure S4, related to Figure 4. Few hits from CD4-ISG screens are donor- or virus-specific.** CD4-ISG screen hits were determined by selecting the genes that scored above background for every screen (56 genes, see Figure 4C). One long non-coding RNA, one previous ISG target (IFITM1), and genes with low expression in CD4$^+$ T cells (32 genes) were removed. MAGeCK scores for the remaining 22 genes are presented here. Graphs include data from all eight screens, separated by virus. Dotted line denotes the highest background threshold from Figure 4B plots. *p<0.05 by Wilcoxon rank test.
