## Supplementary material for "Several cell-intrinsic effectors drive type I interferon-mediated restriction of HIV-1 in primary CD4^+^ T cells": Figure S5

**Figure S5, related to Figure 6.**

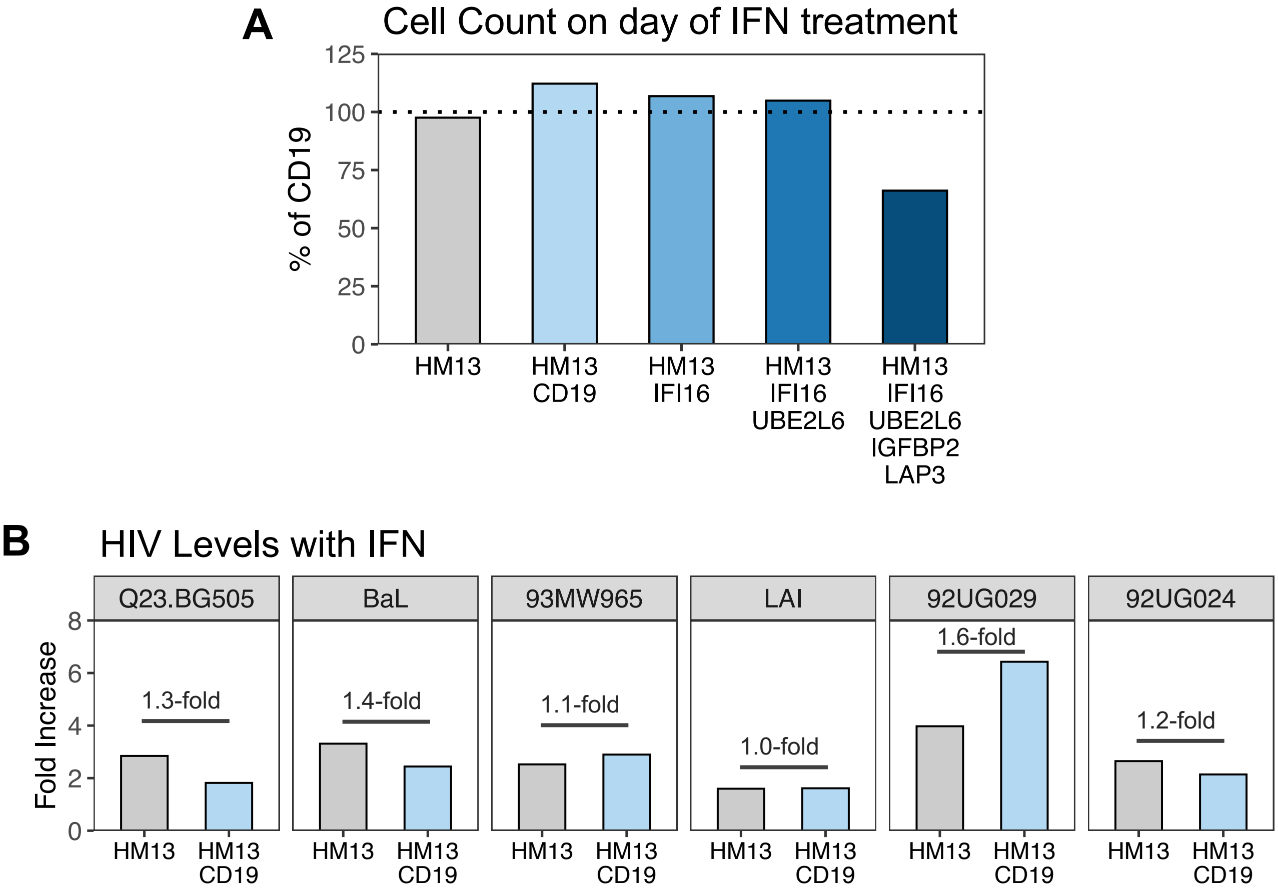


**Figure S5, related to Figure 6. Supporting data for multi-gene knockout experiment.**

(**A**) The number of cells for each ISG KO on the day of IFN treatment are shown relative to CD19-edited cells. Dotted line at 100%. (**B**) The increase in HIV Levels with IFN (ISG KO/CD19 KO) for HM13 single or double KO as compared to CD19 KO. The fold difference between HM13 single and double KO is depicted within each plot.
