## Supplementary material for "Several cell-intrinsic effectors drive type I interferon-mediated restriction of HIV-1 in primary CD4^+^ T cells": Figure S6

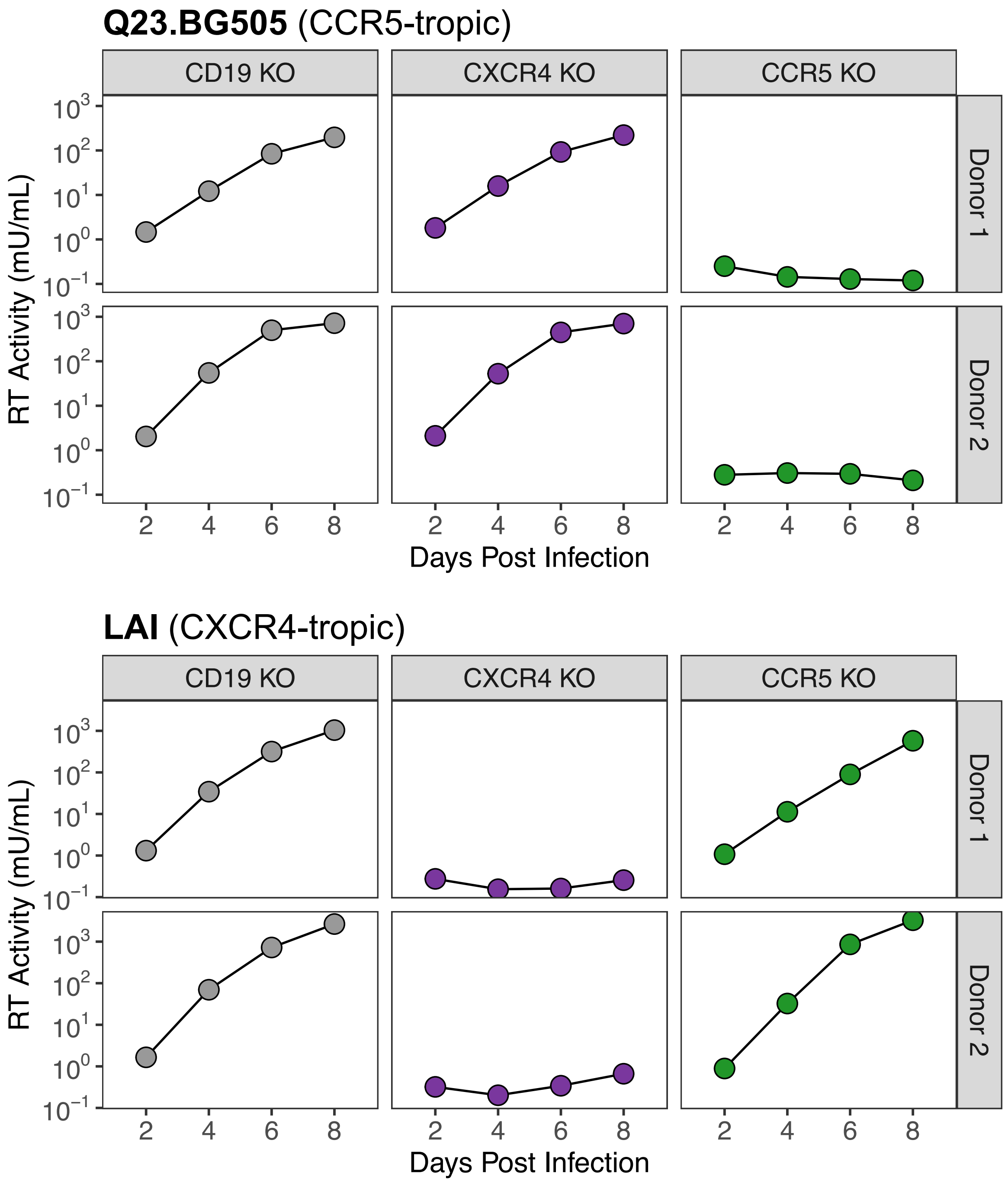


**Figure S6. CD19 editing does not impact HIV infection.**

To determine whether CD19 editing impacts HIV infection, we edited CD19 and HIV co-receptors, CXCR4 and CCR5, in $CD4^+$ T cells from two donors. CXCR4 editing should not impact CCR5-tropic infection (Q23.BG505), and vice-versa for LAI infection. Spreading infection (MOI=0.02) data is shown for all treatments.
